# Novel isolates expand the physiological diversity of *Prochlorococcus* and illuminate its macroevolution

**DOI:** 10.1101/2023.12.03.569780

**Authors:** Jamie W. Becker, Shaul Pollak, Jessie W. Berta-Thompson, Kevin W. Becker, Rogier Braakman, Keven D. Dooley, Thomas Hackl, Allison Coe, Aldo Arellano, Kristen N. LeGault, Paul M. Berube, Steven J. Biller, Andrés Cubillos-Ruiz, Benjamin A. S. Van Mooy, Sallie W. Chisholm

## Abstract

*Prochlorococcus* is a diverse picocyanobacterial genus and the most abundant phototroph on Earth. Its photosynthetic diversity divides it into high- or low-light adapted groups representing broad phylogenetic grades - each composed of several monophyletic clades. Here we physiologically characterize four new *Prochlorococcus* strains isolated from below the deep chlorophyll maximum in the North Pacific Ocean and combine this information with genomic and evolutionary analyses. The isolates belong to deeply-branching low-light adapted clades that have no other cultivated representatives and display some unusual characteristics. For example, despite its otherwise low-light adapted physiological characteristics, strain MIT1223 has low chl *b_2_* content similar to high-light adapted strains. Isolate genomes revealed that each strain contains a unique arsenal of pigment biosynthesis and binding alleles that have been horizontally acquired, contributing to the observed physiological diversity. Comparative genomic analysis of all picocyanobacteria reveals that Pcb, the major pigment carrying protein in *Prochlorococcus*, greatly increased in copy number and diversity per genome along a branch that coincides with the loss of facultative particle attachment. Collectively, these observations add support to the current macroevolutionary model of picocyanobacteria, where niche constructing radiations allowed ancestral lineages to transition from a particle-attached to planktonic lifestyle and broadly colonize the water column, followed by adaptive radiations near the surface that pushed ancestral lineages deeper in the euphotic zone resulting in modern depth-abundance profiles.

**Originality-Significance Statement:** The marine cyanobacterium, *Prochlorococcus*, is among the Earth’s most abundant organisms, and much of its genetic and physiological diversity remains uncharacterized. While field studies help reveal the scope of diversity, cultured isolates allow us to link genomic potential to physiological processes, illuminate eco-evolutionary feedbacks, and test theories arising from comparative genomics of wild cells. Here, we report the isolation and characterization of novel low-light (LL) adapted *Prochlorococcus* strains that fill in multiple evolutionary gaps. These new strains are the first cultivated representatives of the LLVII and LLVIII paraphyletic grades of *Prochlorococcus*, which are broadly distributed in the lower regions of the ocean euphotic zone. Each of these grades is a unique, highly diverse section of the *Prochlorococcus* tree that separates distinct ecological groups: the LLVII grade branches between monophyletic clades that have facultatively particle-associated and constitutively planktonic lifestyles, while the LLVIII grade lies along the branch that leads to all high-light (HL) adapted clades. Characterizing strains and genomes from these grades yields insights into the large-scale evolution of *Prochlorococcus*.

The new LLVII and LLVIII strains are adapted to growth at very low irradiance levels and possess unique light-harvesting gene signatures and pigmentation. The LLVII strains represent the most basal *Prochlorococcus* group with a major expansion in photosynthetic antenna genes. Further, a strain from the LLVIII grade challenges the paradigm that all LL-adapted *Prochlorococcus* exhibit high ratios of chl *b*:*a_2_*. These findings provide insights into major transitions in *Prochlorococcus* evolution, from the benthos to a fully planktonic lifestyle and from growth at low irradiances to the rise of the HL-adapted clades that dominate the modern ocean.

## Introduction

The picocyanobacterium *Prochlorococcus* is numerically the most abundant photoautotrophic organism in the global ocean (Flombaum *et al*., 2013). It represents an enormous and genetically diverse collective of cells (Kashtan *et al*., 2014) tuned to thrive under particular environmental conditions (Kettler *et al*., 2007; Biller *et al*., 2014). *Prochlorococcus* cells have traditionally been divided into high-light (HL) and low-light (LL)-adapted groups based on their optimal light intensity for growth, which generally corresponds to the depths at which they display their maximal abundance (Moore *et al*., 1995, 1998; West and Scanlan, 1999). Finer scale niche partitioning among lineages has been linked to other abiotic factors, including temperature, trace metal and inorganic nitrogen acquisition, and vertical mixing (Bouman *et al*., 2006; Johnson *et al*., 2006; Zinser *et al*., 2007; Thompson *et al*., 2021). Over the years since its discovery (Chisholm *et al*., 1988), environmental sequencing efforts throughout the global ocean have continued to uncover novel genetic diversity within the *Prochlorococcus* collective (Coleman and Chisholm, 2007; Berube *et al*., 2018; Biller, Berube, *et al*., 2018).

Within the broadly defined HL and LL-adapted groups of *Prochlorococcus*, monophyletic clades have been designated based on sequence similarity of the 16S/23S rRNA intergenic spacer sequence (ITS) (Rocap *et al*., 2002; Biller *et al*., 2015). Laboratory isolates of *Prochlorococcus* have existed since shortly after its discovery (Chisholm *et al*., 1992; Partensky *et al*., 1993), and while more than 100 isolates are now available, environmental sequencing continues to reveal groups within the *Prochlorococcus* collective for which no representative isolates exist. These include the HLIII, HLIV, and HLV lineages typically found in high-nutrient, low-chlorophyll environments (Rusch *et al*., 2010; West *et al*., 2010; Huang *et al*., 2011; Malmstrom *et al*., 2012), the HLVI lineage (Huang *et al*., 2011), and the basal LLV/AMZ1, LLVI/AMZII, and AMZIII lineages associated with oxygen minimum zones (Lavin *et al*., 2010; Ulloa *et al*., 2021). Furthermore, quantitative PCR based abundance measurements typically underestimate the total number of *Prochlorococcus* cells *in-situ* when compared to flow cytometry counts, particularly for samples taken at or near the base of the euphotic zone. Because PCR primers are highly specific and are based on the known diversity of *Prochlorococcus*, this suggests a significant fraction of LL-adapted lineages are not represented in culture collections (Ahlgren *et al*., 2006; Zinser *et al*., 2006; Martiny *et al*., 2009). For some of these uncultivated groups, little is known beyond the fact that they exist, precluding a deeper understanding of the evolutionary history of *Prochlorococcus*.

One particularly abundant, deeply branching uncultivated group originally called NC1 (Martiny *et al*., 2009) and later renamed to LLVII (Biller *et al*., 2015) is situated phylogenetically between two broad groups - LLIV and LLII/III - associated with different lifestyles (Figure 1). The transition between these two groups had major implications for the evolution and ecology of ancient *Prochlorococcus.* That is, the more basal LLIV lineage is capable of particle attachment, while the LLII/III clade marks the earliest diverging lineage of fully planktonic cells that characterize the rest of the *Prochlorococcus* tree (Capovilla *et al*., 2023). The LLVII group was first reported in a 2009 clone library study where they comprised as much as 61.7% (average 23.1%) of the *Prochlorococcus* sequences detected in the lower euphotic zone (140 - 160 m) in the North Atlantic and North Pacific Oceans (Martiny *et al*., 2009). This uncultivated group has also been detected at and below 100 m in the Red Sea, where they were the majority (58%) of LL-adapted sequences recovered (Shibl *et al*., 2014) and in the lower euphotic zone of the western Pacific Ocean and South China Sea (Huang *et al*., 2011; Jiao *et al*., 2013). Therefore, the physiology and genomic content of the LLVII lineage are particularly intriguing as they could shed light on niche differentiation among LL-adapted groups and macroevolution of the *Prochlorococcus* collective.

**Figure 1.**
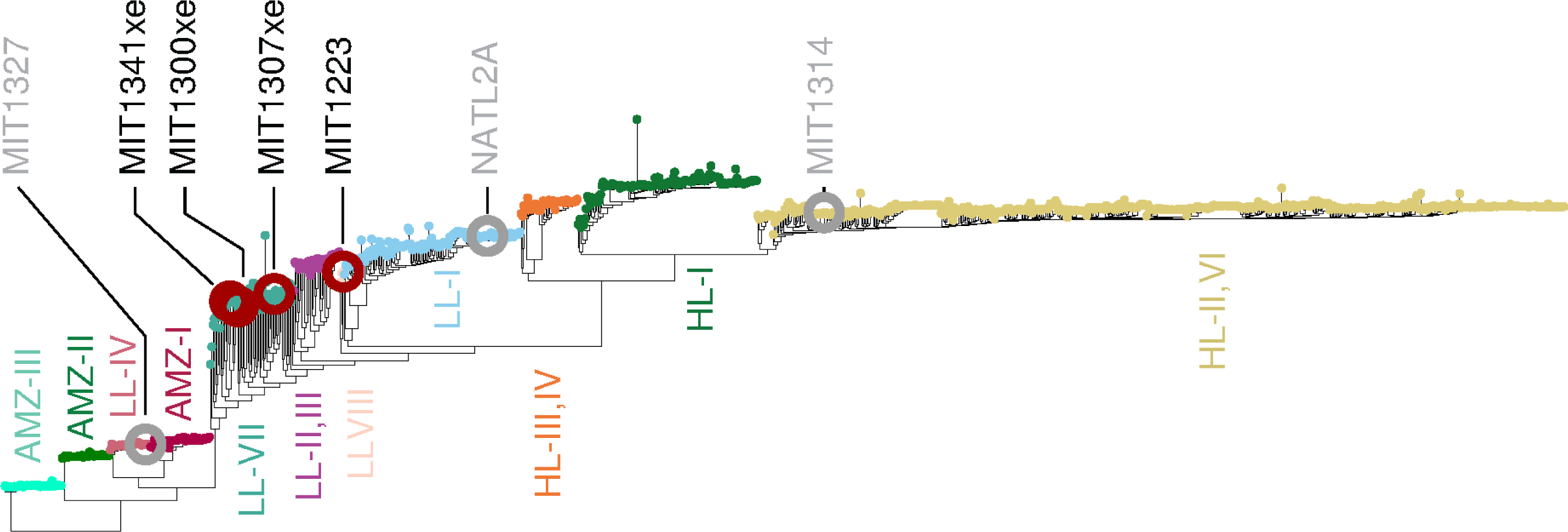
Placement of the isolates used in this study in a phylogeny of 1200 non-identical Prochlorococcus genomes. The genome database contains isolate genomes, MAGs, and SAGs with a wide range of genome completeness, isolation sources, and sequencing methods (see methods). The tree was built using FastTree2 (Price *et al*., 2010) on the concatenated alignment of 44 synteny-conserved orthologous genes (methods). Major clades are represented by tip color. Novel isolates (black text, red circles) and representative strains (gray text and circles) are indicated on the tree.

The current view of this evolutionary history is that *Prochlorococcus* diverged from its sister lineage, *Synechococcus*, ca. 420-300 million years ago (Fournier *et al*., 2021; Capovilla *et al*., 2023). Combining genomic, physiological, metabolic, elemental, and environmental data suggests a macroevolutionary model regarding the diversification of major *Prochlorococcus* lineages (Braakman *et al*., 2017; Berube *et al*., 2019; Capovilla *et al*., 2023) in which the picocyanobacterial ancestor had a mostly particle-attached lifestyle that was lost in the ancestor of all LLVII. We speculate that this LLVII ancestor was committed to a purely planktonic lifestyle, broadly colonizing the water column. Over time adaptive radiations that allowed cells to devote more photosynthetic electron flux to nutrient uptake, effectively drawing surface nutrient levels lower and lower, made surface waters uninhabitable to ancestral lineages (Braakman *et al*., 2017). This resulted in modern ecotype depth-abundance distributions (Johnson *et al*., 2006; Zinser *et al*., 2006; Malmstrom *et al*., 2010; Thompson *et al*., 2021), where more deeply diverging ecotypes are differentially more abundant at greater depths, where photosynthetic electron flux is lower and does not allow full drawdown of nutrient levels (Braakman *et al*., 2017). While understanding deep-time evolution is inherently inferential, studying modern deep-branching lineages has the potential to inform us about the genomic content and physiology of ancient common ancestors, and further constrain possible scenarios.

Here, we study the physiology and genomics of deep-branching *Prochlorococcus* lineages and use their properties to refine our synoptic macroevolutionary perspective. We describe a culturing pipeline explicitly targeting the acquisition of deep-branching low-light adapted lineages, resulting in the isolation of four novel strains belonging to basal lineages. The isolates were physiologically characterized in terms of broad niche-determining traits such as light and temperature optima for growth, photosynthetic capacity, and pigment content. We sequenced their genomes and examined genetic diversity related to pigment biosynthesis and photosynthetic antenna genes of the *pcb* family, which are the primary chlorophyll-binding proteins in *Prochlorococcus*. We conclude by merging the physiological properties and genomic information of these strains with the current macroevolutionary model of forces that drove *Prochlorococcus* diversification in ancient oceans.

## Results and Discussion

### Phylogenetic placement of isolates

Four *Prochlorococcus* strains were isolated from below the deep chlorophyll maximum and 1% light level (Table 1) using cultivation methods designed to enrich LL-adapted cells. The four strains proved difficult to identify using standard sequence alignment of their ITS regions; they had < 70% nucleotide identity to strains from established clades and each other (Figure S1). These poor alignments precluded the generation of ITS-based phylogenetic trees with reliable bootstrap values, thus we moved to sequence alignments generated using 44 marker genes with conserved synteny and compared them with 1200 non-identical *Prochlorococcus* isolate genomes, SAGs, and MAGs. The four isolates belong to two distinct paraphyletic regions lacking any cultivated representatives (Figure 1). One of these regions, encompassing strains MIT1300xe, MIT1307xe and MIT1341xe (where an xe suffix designates xenic cultures, and all other cultures are axenic), falls between the LLII/III and LLIV clades and includes genomes with ITS sequences that resemble those of the NC1 group (later renamed LLVII) discovered in environmental clone libraries (Martiny *et al*., 2009; Biller *et al*., 2015). Our isolates, along with SAGs from this group (Berube *et al*., 2018), confirm hypotheses that these lineages are not monophyletic (Martiny *et al*., 2009; Huang *et al*., 2011). The fourth isolate (MIT1223) resides in a paraphyletic region located between the LLI and LLII/III clades.

**Table 1.** Origin of the *Prochlorococcus* strains described in this work. *Denotes strains representing lineages with no prior isolates. The isolation location for all strains was Station ALOHA in the subtropical North Pacific Ocean (22.75 °N 158 °W). The light column indicates the range of irradiance values experienced by the cells during the journey from enrichment to unialgal isolate and the medium column the nutrient concentrations if they differ from Pro2 and Pro99 media recipes as described in (Moore *et al*., 2007).

| Strain | Clade or Grade | Sampling Date | Sampling Depth (m) | Isolation Medium | Pre-filtration | Purification Method | Light ($\mu\text{mol photons m}^{-2} \text{ s}^{-1}$) | Notes |
| --- | --- | --- | --- | --- | --- | --- | --- | --- |
| MIT1314 | HLII | June 2013 | 150 | Pro2 + 1 $\mu\text{M}$ sodium thiosulfate | Gravity (1.0 $\mu\text{m}$) | dilution to extinction | 0.3 - 40 | |
| MIT1223 | LLVIII* | Sept. 2012 | 175 | 16 $\mu\text{M}$ $\text{NO}_3^-$<br>1 $\mu\text{M}$ $\text{PO}_4^{3-}$<br>1/10<sup>th</sup> Pro99 metals | None | dilution to extinction | 3 - 19 | Additional filtration (0.8 $\mu\text{m}$) |
| MIT1300xe | LLVII* | June 2013 | 150 | 15 $\mu\text{M}$ $\text{NO}_2^-$<br>1 $\mu\text{M}$ $\text{PO}_4^{3-}$<br>Pro2/Pro99 trace metals | Gravity (1.0 $\mu\text{m}$) | non axenic | 0.3 - 12 | |
| MIT1307xe | LLVII* | June 2013 | 150 | 15 $\mu\text{M}$ $\text{NO}_2^-$<br>1 $\mu\text{M}$ $\text{PO}_4^{3-}$<br>Pro2/Pro99 trace metals | Gravity (1.0 $\mu\text{m}$) | non axenic | 0.3 - 12 | Additional filtration (0.8 $\mu\text{m}$) |
| MIT1341xe | LLVII* | June 2013 | 150 | 15 $\mu\text{M}$ $\text{NO}_2^-$<br>1 $\mu\text{M}$ $\text{PO}_4^{3-}$<br>Pro2/Pro99 trace metals | Gravity (1.0 $\mu\text{m}$) | non axenic | 0.3 - 12 | Additional filtration (0.8 $\mu\text{m}$) |
| MIT1327 | LLIV | June 2013 | 150 | Pro2 + 1 $\mu\text{M}$ sodium thiosulfate | Gravity (1.0 $\mu\text{m}$) | dilution to extinction | 3 - 12 | |

The fact that all four of these isolates were obtained from the base of the euphotic zone and maintained under low irradiances in the laboratory, along with their phylogenetic placement with previously identified LL-adapted clades, supports hypotheses that these lineages are LL-adapted (Martiny *et al*., 2009; Shibl *et al*., 2014). Here, we refer to the paraphyletic region between the LLII/III and LLIV clades (encompassing strains MIT1300xe, MIT1307xe and MIT1341xe) as the LLVII grade and the paraphyletic region between the LLI and LLII/III clades (encompassing strain MIT1223) as the LLVIII grade (Figure 1).

To further understand these novel groups, we analyzed pigment content and growth rate effects of light and temperature on one isolate from each grade alongside representative HL and LL-adapted strains from previously described monophyletic clades. Strains MIT1314 (HLII) and MIT1327 (LLIV) were chosen to represent two extremes of the *Prochlorococcus* collective life-history spectrum. MIT1327 (LLIV) represents a deep-branching low-light adapted strategy that can be particle-associated and thrives in the nutrient rich conditions of the lower water-column (Capovilla *et al*., 2023), while MIT1314 (HLII) represents a recently diverging high-light adapted strategy with a streamlined genome that is common in the oligotrophic open ocean (Moore and Chisholm, 1999). To understand how our novel isolates, which are phylogenetically placed between these two representative strains, lie between the two life-history extremes, we focused our experiments on MIT1223 from the LLVIII grade and MIT1300xe as a representative of the LLVII grade.

### Photophysiology

#### Light-dependent growth rates

Growth rates as a function of light intensity were similar for MIT1223, MIT1300xe, and MIT1327 (LLIV clade), but distinct from MIT1314 (HLII clade) - supporting the notion that the paraphyletic LLVII and LLVIII grades comprise LL-adapted lineages (Figure 2A). MIT1223, MIT1300xe, and MIT1327 grew at nearly maximal rates at irradiance levels that were too low to support the growth of MIT1314. Conversely, MIT1314 grew well at the highest irradiance (200 μmol photons m^-2^ s^-1^), while the other strains did not grow reliably at intensities ≥ 73 μmol photons m^-2^ s^-1^ (Table 2A).

**Figure 2.**
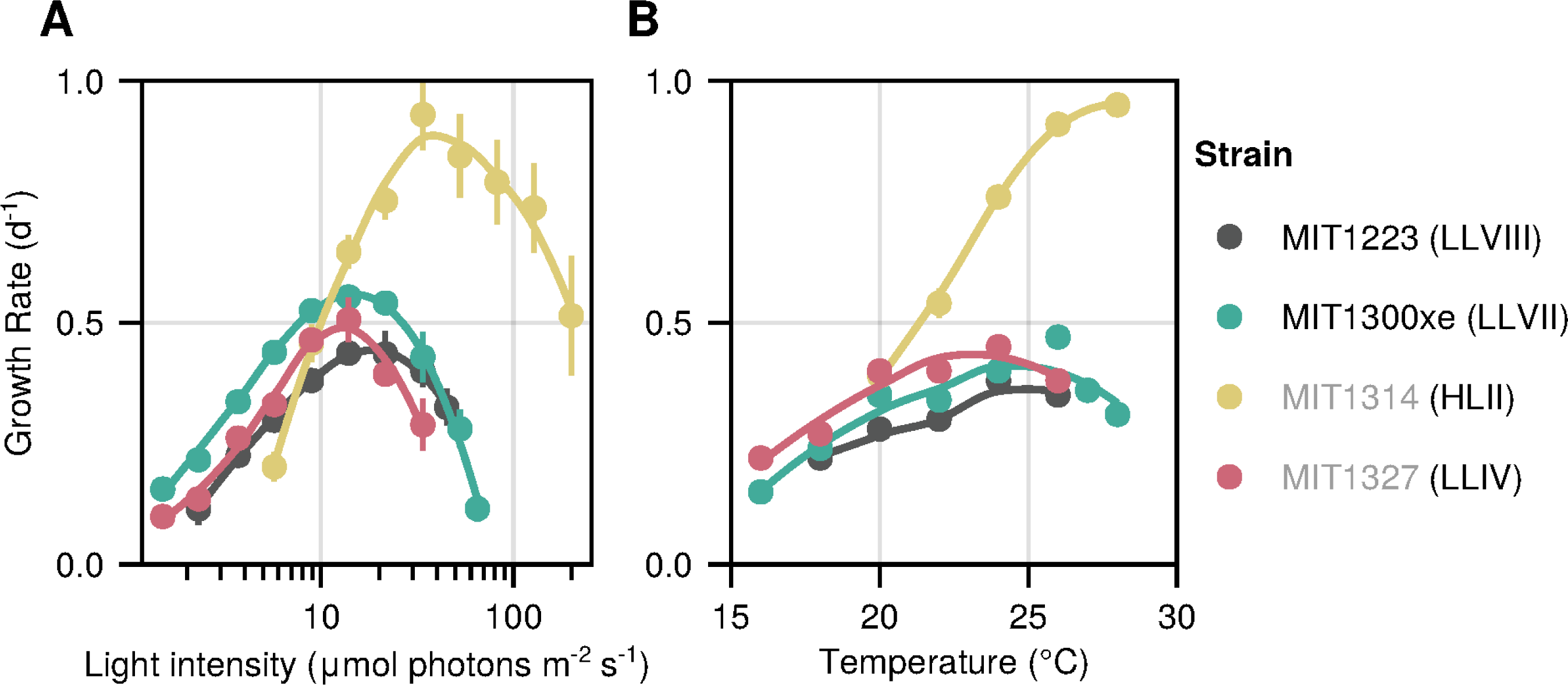
The light and temperature dependent growth rate of novel *Prochlorococcus* isolates. Growth rate as a function of light intensity (A) and temperature (B). Regressions were produced by locally weighted smoothing (LOESS). Circles represent the mean (± s.d.) of biological replicate cultures acclimated to each condition (see Table 2). Experiments in panel A were conducted at 24 ± 1 °C and experiments in panel B were done at 76 ± 1 µmol photons m^-2^ s^-1^ for MIT1314; 20 ± 1 µmol photons m^-2^ s^-1^ for MIT1223, MIT1300xe and MIT1327 on a 14:10 h light:dark cycle. Error bars are smaller than the size of the symbols where not visible. Novel isolate names are marked with black text, while previously described strains are listed in gray. The clade/grade designation of each strain is indicated in parenthesis.

**Table 2.**
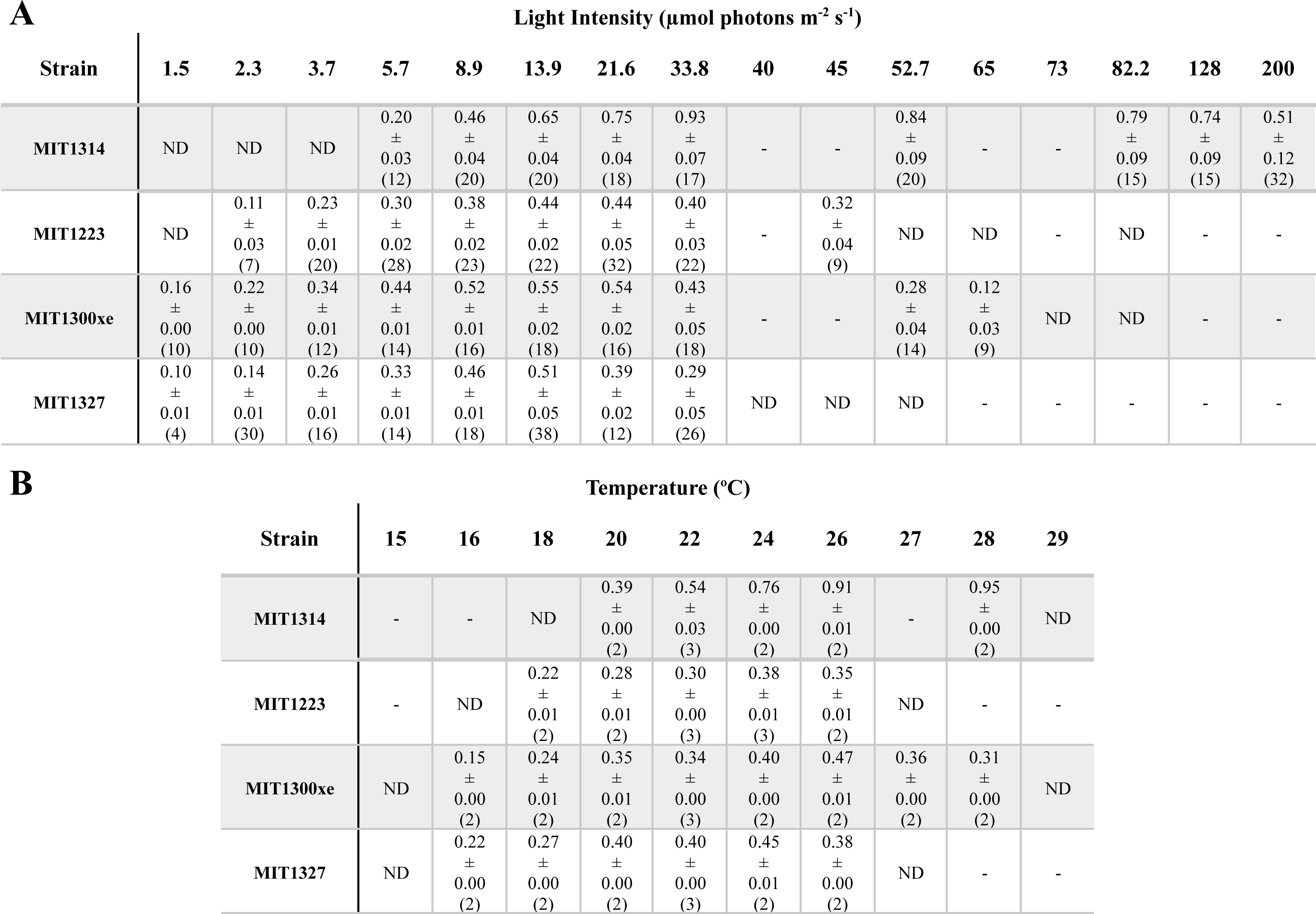
Mean growth rates (d^-1^) ± s.d. for *Prochlorococcus* strains acclimated to different light intensities (A) and temperatures (B). ND (not determined) indicates that the strain was tested at this condition, however a consistent reproducible growth rate could not be achieved. A dash indicates that growth at this condition was not tested. Parentheses indicate the number of biological replicates included in calculations.

**A**
| Strain | Light Intensity ($\mu\text{mol photons m}^{-2} \text{ s}^{-1}$) | | | | | | | | | | | | | | | |
| --- | --- | --- | --- | --- | --- | --- | --- | --- | --- | --- | --- | --- | --- | --- | --- | --- |
|  | 1.5 | 2.3 | 3.7 | 5.7 | 8.9 | 13.9 | 21.6 | 33.8 | 40 | 45 | 52.7 | 65 | 73 | 82.2 | 128 | 200 |
| MIT1314 | ND | ND | ND | 0.20<br>±<br>0.03<br>(12) | 0.46<br>±<br>0.04<br>(20) | 0.65<br>±<br>0.04<br>(20) | 0.75<br>±<br>0.04<br>(18) | 0.93<br>±<br>0.07<br>(17) | - | - | 0.84<br>±<br>0.09<br>(20) | - | - | 0.79<br>±<br>0.09<br>(15) | 0.74<br>±<br>0.09<br>(15) | 0.51<br>±<br>0.12<br>(32) |
| MIT1223 | ND | 0.11<br>±<br>0.03<br>(7) | 0.23<br>±<br>0.01<br>(20) | 0.30<br>±<br>0.02<br>(28) | 0.38<br>±<br>0.02<br>(23) | 0.44<br>±<br>0.02<br>(22) | 0.44<br>±<br>0.05<br>(32) | 0.40<br>±<br>0.03<br>(22) | - | 0.32<br>±<br>0.04<br>(9) | ND | ND | - | ND | - | - |
| MIT1300xe | 0.16<br>±<br>0.00<br>(10) | 0.22<br>±<br>0.00<br>(10) | 0.34<br>±<br>0.01<br>(12) | 0.44<br>±<br>0.01<br>(14) | 0.52<br>±<br>0.01<br>(16) | 0.55<br>±<br>0.02<br>(18) | 0.54<br>±<br>0.02<br>(16) | 0.43<br>±<br>0.05<br>(18) | - | - | 0.28<br>±<br>0.04<br>(14) | 0.12<br>±<br>0.03<br>(9) | ND | ND | - | - |
| MIT1327 | 0.10<br>±<br>0.01<br>(4) | 0.14<br>±<br>0.01<br>(30) | 0.26<br>±<br>0.01<br>(16) | 0.33<br>±<br>0.01<br>(14) | 0.46<br>±<br>0.01<br>(18) | 0.51<br>±<br>0.05<br>(38) | 0.39<br>±<br>0.02<br>(12) | 0.29<br>±<br>0.05<br>(26) | ND | ND | ND | - | - | - | - | - |

**Table 2.** Mean growth rates ( $\text{d}^{-1}$ ) ± s.d.
| Strain | Temperature ($^{\circ}\text{C}$) | | | | | | | | | |
| --- | --- | --- | --- | --- | --- | --- | --- | --- | --- | --- |
|  | 15 | 16 | 18 | 20 | 22 | 24 | 26 | 27 | 28 | 29 |
| MIT1314 | - | - | ND | 0.39<br>±<br>0.00<br>(2) | 0.54<br>±<br>0.03<br>(3) | 0.76<br>±<br>0.00<br>(2) | 0.91<br>±<br>0.01<br>(2) | - | 0.95<br>±<br>0.00<br>(2) | ND |
| MIT1223 | - | ND | 0.22<br>±<br>0.01<br>(2) | 0.28<br>±<br>0.01<br>(2) | 0.30<br>±<br>0.00<br>(3) | 0.38<br>±<br>0.01<br>(3) | 0.35<br>±<br>0.01<br>(2) | ND | - | - |
| MIT1300xe | ND | 0.15<br>±<br>0.00<br>(2) | 0.24<br>±<br>0.01<br>(2) | 0.35<br>±<br>0.01<br>(2) | 0.34<br>±<br>0.00<br>(3) | 0.40<br>±<br>0.00<br>(2) | 0.47<br>±<br>0.01<br>(2) | 0.36<br>±<br>0.00<br>(2) | 0.31<br>±<br>0.00<br>(2) | ND |
| MIT1327 | ND | 0.22<br>±<br>0.00<br>(2) | 0.27<br>±<br>0.00<br>(2) | 0.40<br>±<br>0.00<br>(2) | 0.40<br>±<br>0.00<br>(3) | 0.45<br>±<br>0.01<br>(2) | 0.38<br>±<br>0.00<br>(2) | ND | - | - |

MIT1300xe (LLVII grade) sustained growth at extremely low light intensities, with a growth rate 1.6 times faster than MIT1327 (LLIV clade) - the only other strain capable of growth at 1.5 μmol photons m^-2^ s^-1^ - reflecting MIT1300xe’s ability to thrive at the base of the euphotic zone where light limits primary production. MIT1300xe also grew well over a broad range of light intensities (1.5 to 65 μmol photons m^-2^ s^-1^) consistently faster than the other LL-adapted strains (Figure 2A). We note, however, that MIT1300xe was the only xenic strain tested and we cannot rule out heterotrophic cells influencing these results due to *Prochlorococcus* mixotrophy (Wu *et al*., 2022) and reduction of oxidative stress (Morris *et al*., 2008; Sher *et al*., 2011; Coe *et al*., 2016; Ma *et al*., 2017). Both MIT1300xe and MIT1223 grew better at higher light intensities than MIT1327, suggesting that members of the LLVII and LLVIII grades may tolerate higher irradiances than those of the LLIV clade. Interestingly, despite originating from depths that typically experience < 5 μmol photons m^-2^ s^-1^, all four strains achieved their maximum growth rates at similar light intensities (13.9 - 33.8 μmol photons m^-2^ s^-1^). However, the maximal growth rate of MIT1314 (HLII clade) was 1.7 times faster than the maximal growth rate of any LL-adapted strain (Table 2A). These results reinforce the well-established notion that adaptation to high-light conditions probably occurred only once and relatively late during *Prochlorococcus* evolution (Kettler *et al*., 2007).

#### Photosynthetic properties

Fast repetition rate fluorometry was used to compare photosynthetic efficiencies and the functional size of light-harvesting antennae among strains when grown at various irradiances. Photosynthetic quantum efficiency (*F*_V_/*F*_M_) generally decreased with increasing light intensity in all four strains, with MIT1327 (LLIV clade) exhibiting the steepest decline, especially at light intensities > 8.9 μmol photons m^-2^ s^-1^ (Figure 3A). MIT1300xe (LLVII grade) had the same photosynthetic quantum efficiency when grown at 1.5 to 21.6 μmol photons m^-2^ s^-1^, but declined at higher intensities, which is in agreement with its consistently high growth rate over this irradiance range when compared to other LL-adapted strains. Interestingly, MIT1300xe is the only strain tested that displayed maximum photosynthetic efficiency when growing at its maximum rate, likely due to the presence of heterotrophs reducing oxidative stress. MIT1327 (LLIV clade) was the most photosynthetic efficient strain at low light intensities, while MIT1300xe (LLVII grade) was the most photosynthetic efficient strain at the intermediate light intensities. Photosystem II (PSII) functional absorption cross-sections (σPSII) were largest at intermediate light intensities, with a general decline in σPSII values for all strains at light intensities greater than those at which the maximum growth rate was observed, potentially indicating light-induced damage to PSII at higher irradiances (Figure 3B). MIT1300xe had larger σPSII values than the other LL-adapted strains at all light intensities, an advantage that is consistent with its faster growth rates.

**Figure 3.**
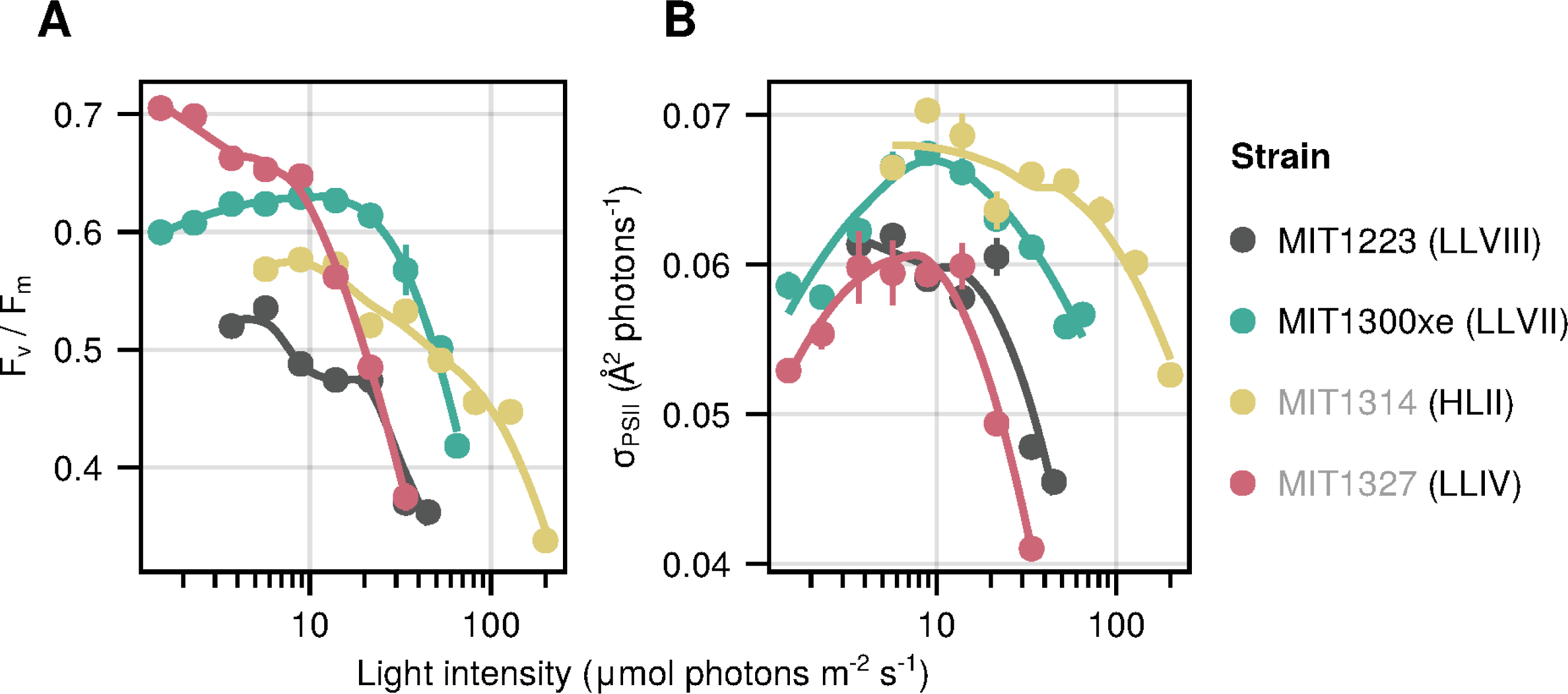
Photosynthetic parameters of novel isolates as a function of growth light intensity. Photosynthetic quantum efficiency (A) and functional absorption cross-section (σPSII) (B). Regressions produced by locally weighted smoothing (LOESS). Circles represent the mean (± s.d.) of biological replicate cultures acclimated to each condition sampled over time (n = 6 to 10). Error bars are smaller than the size of the symbols where not visible. Novel isolate names are marked with black text, while previously described strains are listed in gray. The clade/grade designation of each strain is indicated in parenthesis.

The photosynthetic properties of MIT1300xe (LLVII) place it as an intermediate in the *Prochlorococcus* collective life-history spectrum, consistent with its phylogenetic placement between the LLIV clade and other ecotypes (Figure 1). MIT1300xe exhibited the greatest photosynthetic quantum efficiency at intermediate light levels (Figure 3A) and σPSII values between those of the LLIV and HLII strains at all light intensities (Figure 3B). This suggests that the photosynthetic apparatus of MIT1300xe may represent a transitional state between LLIV and later diverging lineages, or that it occupies a niche where these intermediate photosynthetic parameters are advantageous. MIT1223ax (LLVIII) however, exhibited low photosynthetic quantum efficiencies despite σPSII values similar to the LLIV strain, suggesting physiological tradeoffs that limit photosynthetic growth efficiency may be a requirement for *Prochlorococcus* cells with with a relatively small genome size and low GC content (Table 3; (Biller *et al*., 2014)) to grow at low irradiances (Figure 2A).

**Table 3.** Genome characteristics and assembly statistics for *Prochlorococcus* strains in this study. *N50 for this assembly was 328.4 kbp.

| Strain | Clade or Grade | Assembly Size (bp) | % GC | # Scaffolds | Average Coverage | # Coding Sequences | JGI Genome ID | NCBI BioSample # |
| --- | --- | --- | --- | --- | --- | --- | --- | --- |
| MIT1314 | HLII | 1,704,447 | 31.2 | 1 | 73 | 1,926 | 2681813573 | SAMN38315734 |
| MIT1223 | LLVIII | 1,795,922 | 35.7 | 1 | 236 | 1,939 | 2681813568 | SAMN38315731 |
| MIT1300xe | LLVII | 1,855,146 | 41.4 | 1 | 186 | 1,965 | 2681813570 | SAMN38315732 |
| MIT1307xe | LLVII | 2,032,419 | 39.9 | 1 | 317 | 2,140 | 2681813572 | SAMN38315733 |
| MIT1341xe | LLVII | 1,937,096 | 40.1 | 1 | 335 | 2,037 | 2681813574 | SAMN38315735 |
| MIT1327 | LLIV | 2,587,389 | 50.3 | 29* | 204 | 2,642 | 2681812949 | SAMN04490364 |

#### Pigment-driven photoacclimation

*Prochlorococcus* cells tune their pigment content to irradiance (Partensky *et al*., 1993; Moore *et al*., 1995), often revealed as an anticorrelation between chlorophyll (red) fluorescence per cell and growth irradiance in flow cytometric signatures. HL and LL-adapted strains are known to display different slopes, where pigment fluorescence decreases more drastically for LL-adapted cells with increasing light intensity (Moore *et al*., 1995; Moore and Chisholm, 1999). We found this slope to be intermediate for MIT1300xe (LLVII grade), which is similar to previous observations of cells from the LLII/III clade (Figure 4A, see also Moore and Chisholm, 1999). Surprisingly, despite being adapted to grow at low light intensities, MIT1223 (LLVIII grade) appeared similar to MIT1314 (HLII clade) in this regard (Figure 4A). In terms of yellow fluorescence, which is mainly mediated by phycoerythrin in *Prochlorococcus*, the HLII strain (MIT1314), which lacks phycoerythrin biosynthetic potential (Hess *et al*., 1999), represents the lowest extreme, while the LLIV strain (MIT1327) represents the highest. The novel LLVII and LLVIII isolates are intermediate in terms of slope and total yellow fluorescence/cell levels, recapitulating their evolutionary location between the basal LLIV and the more recently emerged HL-adapted clades (Figure 4B).

**Figure 4.**
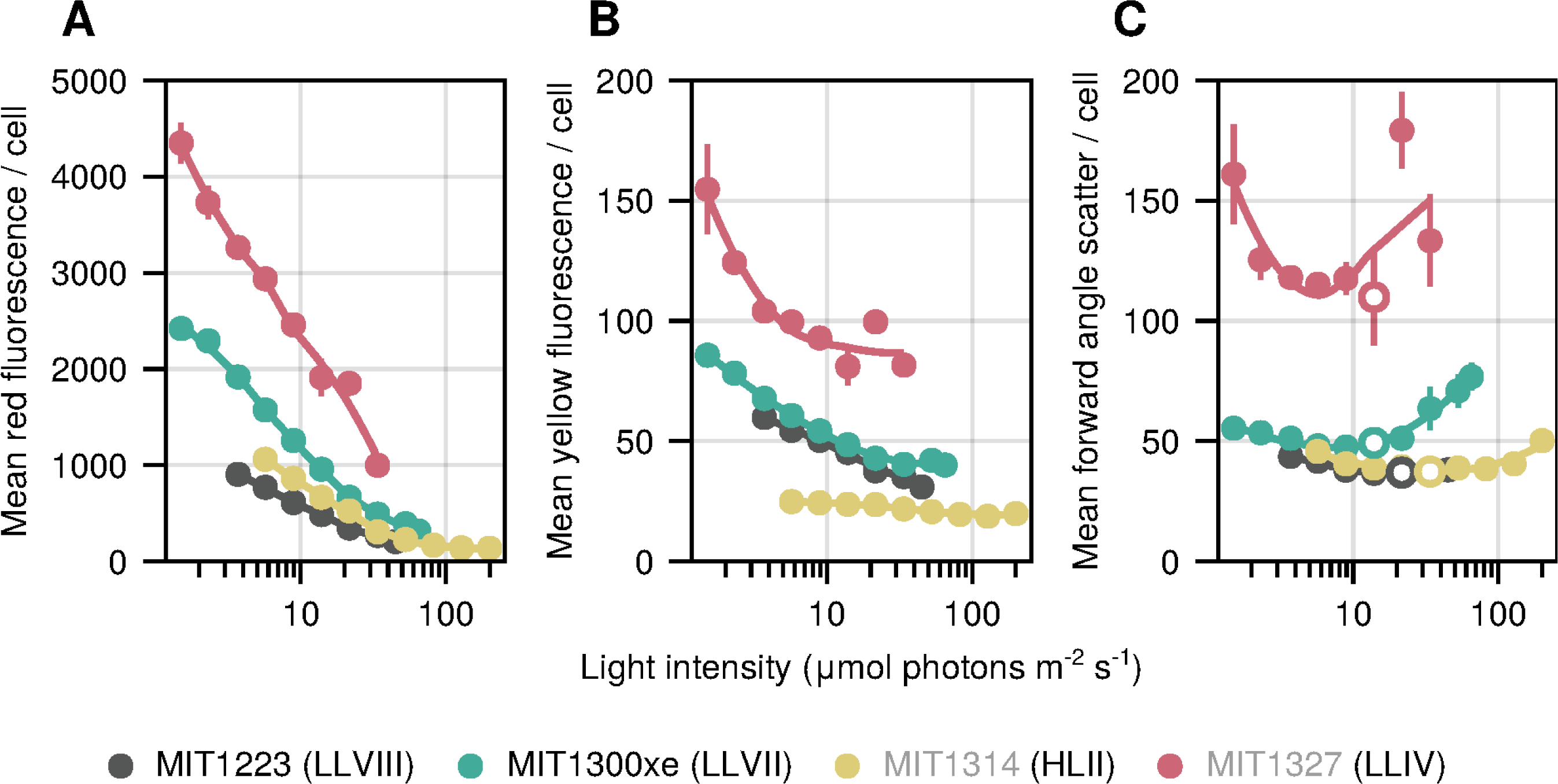
Fluorescence and light scattering properties of novel isolates as a function of growth light intensity. Flow cytometric analyses (A) red (chlorophyll) and (B) yellow (phycoerythrin) fluorescence per cell, and (C) forward light scatter (proxy for cell size). Symbols represent the mean (± s.d.) of biological replicate cultures acclimated to each condition. Error bars are smaller than the size of the symbols where not visible. Open circles in (C) denote the light level at which maximum growth rate was achieved for each strain. Novel isolate names are marked with black text, while representative strains are listed in gray. The lineage of each strain is indicated in parenthesis

All four strains tended to increase in size with increasing irradiance (Figure 4C). As expected, based on their clade and genome size, MIT1327 (LLIV clade) cells were the largest cells at all light intensities (Cermak *et al*., 2016). The forward angle scatter per cell of MIT1223 (LLVIII grade) was similar to that of the HLII strain at all irradiance levels (Figure 4C), providing further evidence that MIT1223 does not follow all canonical descriptions of a LL-adapted *Prochlorococcus* cell. These results provide potential clues about the period of evolutionary tinkering that occurred during the divergence of LLIV and other *Prochlorococcus* - suggesting that changes to the physiology and pigments that occurred as cells adapted to a planktonic lifestyle may have occurred in distinct stages.

To explore whether or not the unexpected properties of MIT1223’s photophysiology - i.e., chlorophyll (red) fluorescence and forward angle scatter per cell - were unique to the LLVIII grade, we examined *Prochlorococcus* strain NATL2A from the LLI clade (Moore *et al*., 1998; Berube *et al*., 2014; Biller *et al*., 2023) acclimated to 4 irradiance levels that fall within the middle of the range tested for the other strains. NATL2A grew faster than the other LL-adapted strains (MIT1223 and MIT1327) at higher light intensities (≥ 26.3) but slower than the HL-adapted strain (MIT1314) (Figure S2A). This finding supports the notion that LLI clade *Prochlorococcus* can tolerate higher irradiance levels than other LL-adapted strains - perhaps due in part to the presence of a photolyase gene and an abundance of high light-inducible genes (Coleman and Chisholm, 2007; Malmstrom *et al*., 2010). Despite its unique light-dependent growth rates (Figure S2A), the *in-vivo* absorption spectra, median chlorophyll (red) fluorescence per cell, and median forward angle scatter per cell of NATL2A (LLI clade) were quite similar to those of MIT1223 (LLVIII grade) and MIT1314 (HLII clade), and distinct from MIT1327 (LLIV clade) (Figure S2 B,C,E). The median yellow (primarily phycoerythrin) fluorescence per cell signature of NATL2A was lower than MIT1223 and similar to that of MIT1314 (Figure S2D). While a detailed pigment analysis of *Prochlorococcus* strains from the LLI clade is needed to confirm these absorption and flow cytometry results, our experiments indicate that despite its phylogenetic placement among LL-adapted clades, NATL2A, like MIT1223, possesses many physiological characteristics that are traditionally associated with HL-adapted strains (e.g. lower red and yellow fluorescence per cell, reduced forward angle light scatter, and likely lower chl *b*:*a_2_*ratios as indicated by *in-vivo* absorption spectra).

*In-vivo* absorption spectra provide further evidence that the pigmentation of MIT1223 is unique among the LL-adapted strains. *Prochlorococcus* cells have traditionally been divided into two major groups based on chlorophyll content with HL-adapted cells exhibiting low total chl *b* (chl *b_1_* + chl *b_2_*) to chl *a_2_* ratios and LL-adapted cells exhibiting high chl *b*:*a_2_* ratios (Partensky *et al*., 1993; Moore *et al*., 1998; Moore and Chisholm, 1999). Comparing spectra for all strains grown at the same low light intensity the second peak at ca. 480 nm (chlorophyll *b_2_*) is visible for MIT1327 (LLIV) and MIT1300xe (LLVII), with higher absorbance than the peak at ca. 450 nm (chlorophyll *a_2_*) (Figure 5). In contrast, the chl *b_2_*peak for MIT1314 (HLII) is lower than its chl *a_2_* peak, matching the canonical definition of HL-adapted strains (Partensky *et al*., 1993; Moore *et al*., 1998). The absorption spectra of MIT1223 (LLVIII) is anomalous for a LL-adapted strain, with no discernible chl *b_2_* peak and a faint peak at ca. 470 nm - likely due to a combination of zeaxanthin and α/β-carotene (Moore *et al*., 1995) - providing further support for atypical pigmentation in the LL-adapted MIT1223 strain.

**Figure 5.**
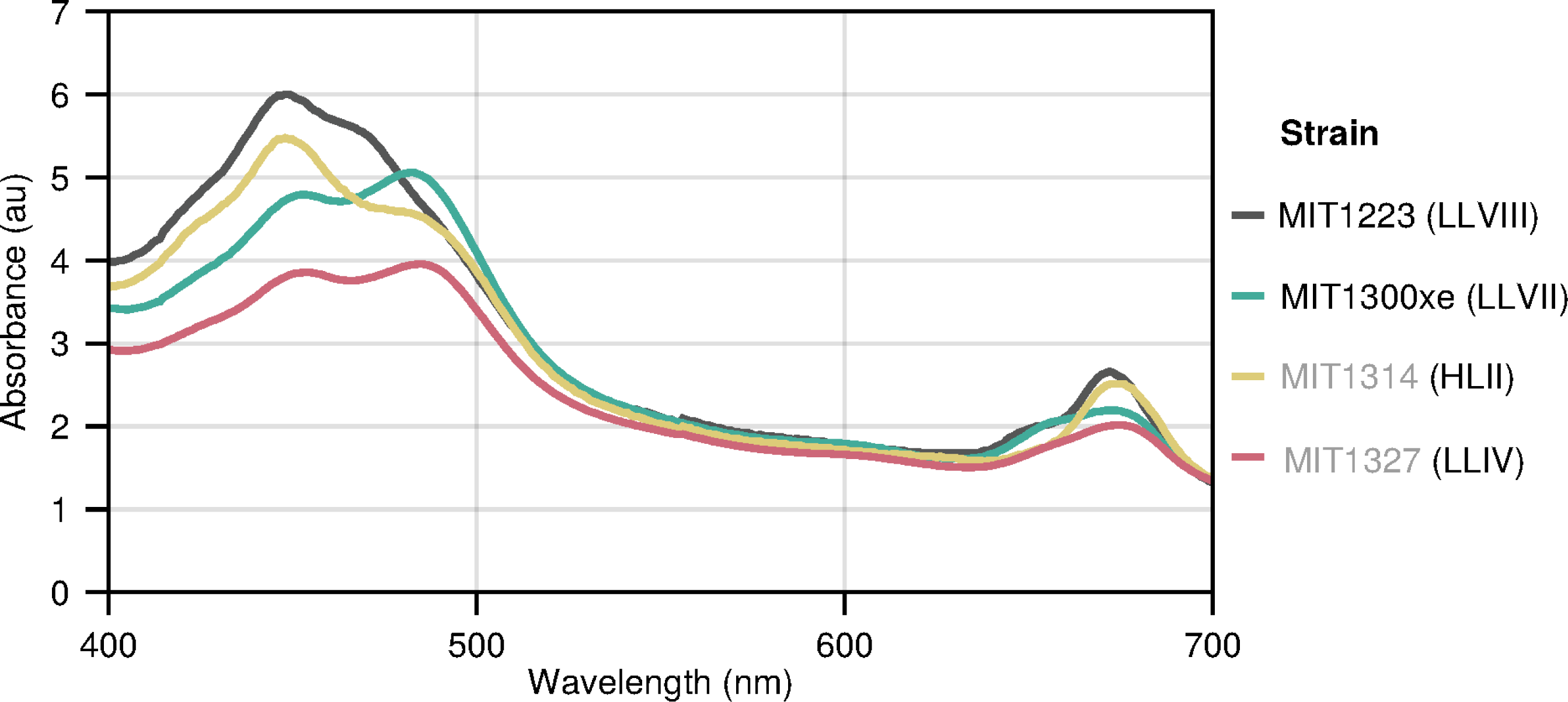
*In vivo* absorption spectra of novel isolates compared to representatives from established LL and HL-adapted clades. Spectra measured for log phase cultures growing at 5.7 μmol photons m^-2^ s^-1^. Novel isolate names are marked with black text, while representative strains are listed in gray. The lineage of each strain is indicated in parenthesis.

We next analyzed pigments using HPLC and MS, which confirmed the *in-vivo* absorption spectra and provided additional information (Figures S3, S4). Strikingly, the chl *b_2_* values for MIT1223 were the lowest of the four strains, including HL-adapted MIT1314 (Figure 6A). LL-adapted strains MIT1327 (LLIV) and MIT1300xe (LLVII) contained appreciable amounts of chl *b_2_*relative to the total measurable pigment content (4 - 39%), especially at lower light intensities, while the relative abundance of chl *b_2_* in MIT1314 (HLII) was lower at all light intensities where both strains could grow (5 - 16%; Figure 6A).

**Figure 6.**
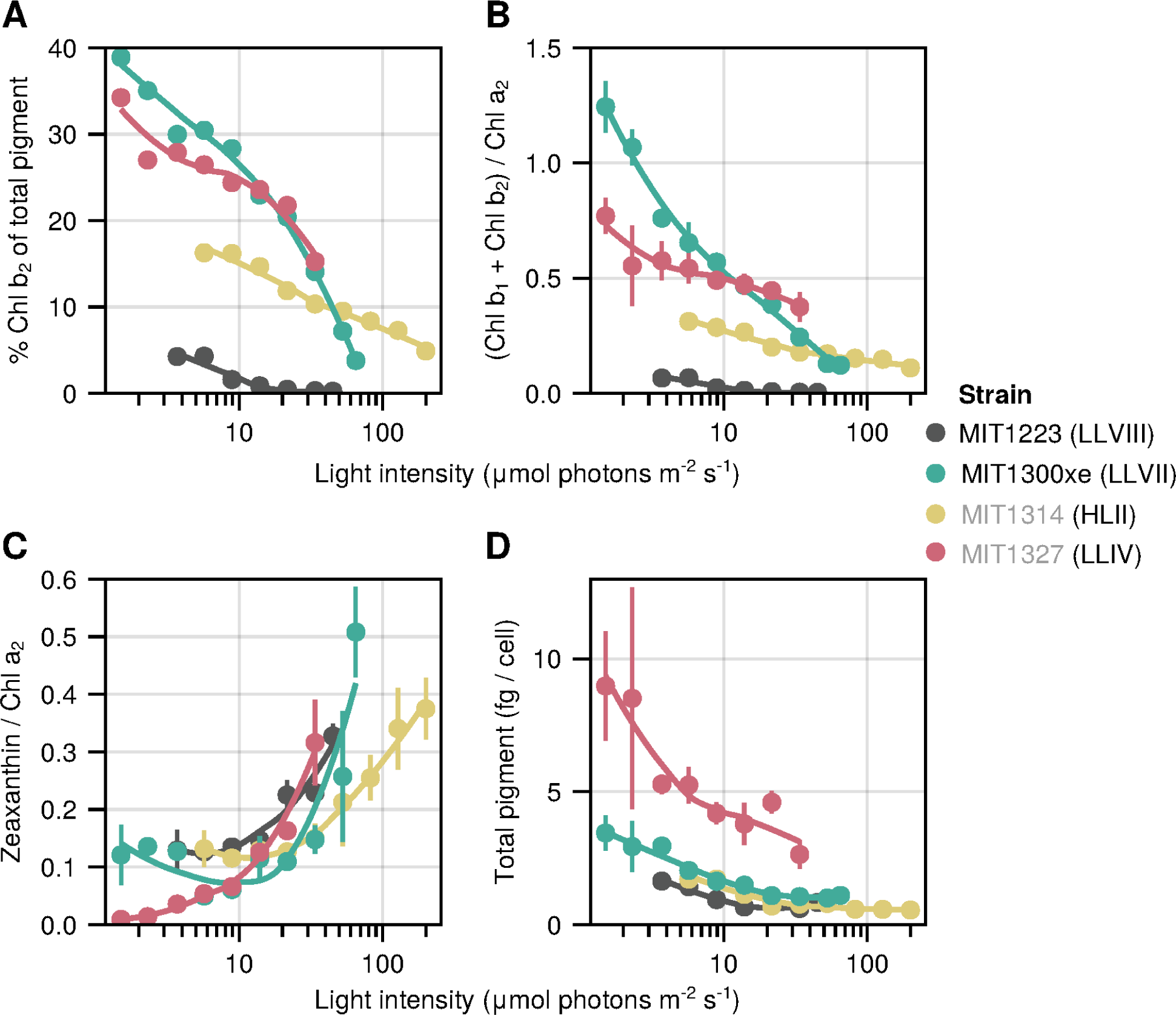
Chlorophyll content of the novel isolates as a function of growth light intensity. Percent of chlorophyll b2 from the total extractable pigment (A), chlorophyll b/a2 ratios (B), zeaxanthin/a2 ratios (C), and total extractable pigment per cell (D). Values are means (± s.d.) of biological replicate cultures acclimated to each condition sampled over time (n = 2 to 7). Lines represent local Loess fits. Novel isolate names are marked with black text, while representative strains are listed in gray. The lineage of each strain is indicated in parenthesis

As expected, the ratio of chl *b*:*a_2_* decreased with increasing light intensity for all strains. MIT1300xe (LLVII) showed ratios > 1 when grown at low intensities (Figure 6B), similar to previous findings for strains from the LLII and LLIV clades (Partensky *et al*., 1993; Moore and Chisholm, 1999; Partensky *et al*., 1999). These high ratios likely facilitate growth at very low irradiances (Moore and Chisholm, 1999). However, the strains tested here still do not have ratios as high as those reported for wild *Prochlorococcus* cells (Goericke and Repeta, 1993), thus the influence of complex heterotrophic partnerships on this ratio may be at play here; the highest chl *b*:*a_2_* ratios *in-vitro* have been measured for xenic cultures (Moore *et al*., 1995; Moore and Chisholm, 1999; this study). MIT1300xe also demonstrated a remarkable capacity for photoacclimation with a range of chl *b*:*a_2_* ratios spanning more than one order of magnitude (Figure 6B). MIT1327 (LLIV) had the next highest chl *b*:*a_2_* ratios; however, these ratios were < 1, even when growing under the lowest intensities (Figure 6B). As expected from previous studies, MIT1314 (HLII) had lower chl *b*:*a_2_* ratios than MIT1300xe and MIT1327 at all light levels with less photoacclimation over the range of light levels tested (Figure 6B). The chl *b*:*a_2_* ratios of MIT1223 (LLVIII) were the lowest of any strain tested - as much as 80 fold lower than MIT1300xe at the same light level. MIT1223 displayed little photoacclimation, and chl *b*:*a_2_* ratios for this strain dropped below 0.01 at the highest light intensities (Figure 6B). MIT1223 is therefore an exception to the long-standing paradigm that LL-adapted strains contain higher ratios of chl *b*:*a_2_* than HL-adapted strains.

Ratios of zeaxanthin to chl *a_2_* generally increased with increasing light intensity, supporting its photoprotective role (Partensky *et al*., 1993; Moore *et al*., 1995) (Figure 6C). MIT1327 had the highest total extractable pigment per cell (2.6 - 9.0 fg/cell) compared to the other three strains tested and MIT1223 had values at or below those for MIT1314 when grown at the same irradiance (Figure 6D). Other primary pigments detected in all four strains consisted of *α/β*-carotene (indistinguishable with our method), pheophytin *a_2_*, Mg 3,8 divinyl pheoporphyrin (DVP) *a_5_*, and an unknown carotenoid (Figures S3, S4, S5), all of which are consistent with previous reports analyzing the pigment content of *Prochlorococcus* cells (Goericke and Repeta, 1992; Partensky *et al*., 1993). Based on its exact mass, the unknown carotenoid has a sum formula of C_40_H_56_O, which matches with both cryptoxanthin and *β*-carotene 5,6-epoxide, however, the MS^2^ fragment data does not appear to match published spectra for either compound (Figure S5). It has been previously suggested that *Prochlorococcus* may contain cryptoxanthin (Goericke and Repeta, 1992), but due to the MS^2^ results we obtained, we do not feel comfortable making a definitive identification, and additional information such as nuclear magnetic resonance analysis is needed to elucidate the structure of this carotenoid. We also putatively identified the compound all-*trans*-retinal in all four strains due to an exact match (retention time and mass) to a standard (Figure S6). However, this identification remains ambiguous, given that the abundance was low, and therefore MS^2^ spectra could not be obtained. We feel the putative identification of this compound is worth noting, however, as the potential ability of *Prochlorococcus* to produce all-*trans*-retinal could have important implications for syntrophic interactions in the marine environment, given the importance of this molecule for proteorhodopsin-based phototrophy (DeLong and Béjà, 2010; Gómez-Consarnau *et al*., 2019).

### Thermophysiology

Temperature-dependent growth rates also revealed similar trends for MIT1223, MIT1300xe, and MIT1327 - which were quite distinct from that of MIT1314 - suggesting the paraphyletic LLVII and LLVIII regions comprise lineages of *Prochlorococcus* that are adapted to colder conditions found deeper in the water column near the base of the euphotic zone - the waters from which they were isolated (Figure 2B). While all of the LL-adapted strains grew much slower than the HL-adapted strain at temperatures ≥ 24 °C, MIT1300xe was still able to grow at the highest temperature at which the HL-adapted strain was able to grow, 2 °C warmer than the upper limit of the other LL-adapted strains (Table 2B). Again, we cannot rule out the potential influence of heterotrophic bacterioplankton, but note that MIT1327 (LLIV clade) grew faster than MIT1300xe at temperatures ≤ 24 °C, despite being axenic. Consistent with the light physiology experiments, MIT1223 generally grew slower than the other strains regardless of temperature, but it could tolerate colder conditions than the HL-adapted strain (Table 2B).

### Photosynthetic antenna evolution

Genomes of the four isolates ranged from 1.8 - 2.0 Mb in size with 35.7 - 41.4 %GC, intermediate to HL-adapted strains and those from the LLIV clade (Biller *et al*., 2014) (Table 3). We examined light-harvesting genes to place these new genomes in context and explore a genetic basis for their observed physiological properties. Unlike other cyanobacteria, most *Prochlorococcus* ecotypes do not produce phycobilisomes as light-harvesting antennae, and instead rely on pigment-binding *pcb* proteins to channel photons to their photosystem II core (Roche *et al*., 1996). This property gives *Prochlorococcus* its distinct fluorescence and absorption profiles and is one of the hallmark adaptations that allowed this group to colonize the entire photic zone (Ting *et al*., 2002). Because these antenna proteins are the major pigment-containing elements in their photosystems, a plausible hypothesis is that the photophysiological features of these novel LLVII/VIII isolates are due – at least in part – to unique combinations of pigment and antenna. Thus, we focused our genomic comparisons on these features.

In agreement with previous reports (Ulloa *et al*., 2021), our analysis shows that most picocyanobacteria encode the iron-inducible *isiA* gene, and *Synecococcus* and the basal *Prochlorococcus* groups (AMZII and AMZIII) encode the full suite of phycobilisome genes, but no *pcb* genes. *pcbA, t*he first *pcb* gene to diverge from *isiA*, first appears in the ancestor of the LLIV/AMZ-I radiation (Figure 7C). A comparison of the genomes from 1,691 *Synechococcus* and *Prochlorococcus* picocyanobacteria (Methods) reveals a major expansion in *pcb* gene copy number and diversity for the LLVII grade that is conserved in later branching LL-adapted clades, but lost in the ancestor of all HL-adapted cells, which only contain *pcbA* (Figure 7). The different ecotypes of *Prochlorococcus* have distinct repertoires of antenna genes, suggesting the observed genetic changes occurred in the ancestor of the group, and that selection has been strong enough to retain these changes.

**Figure 7.**
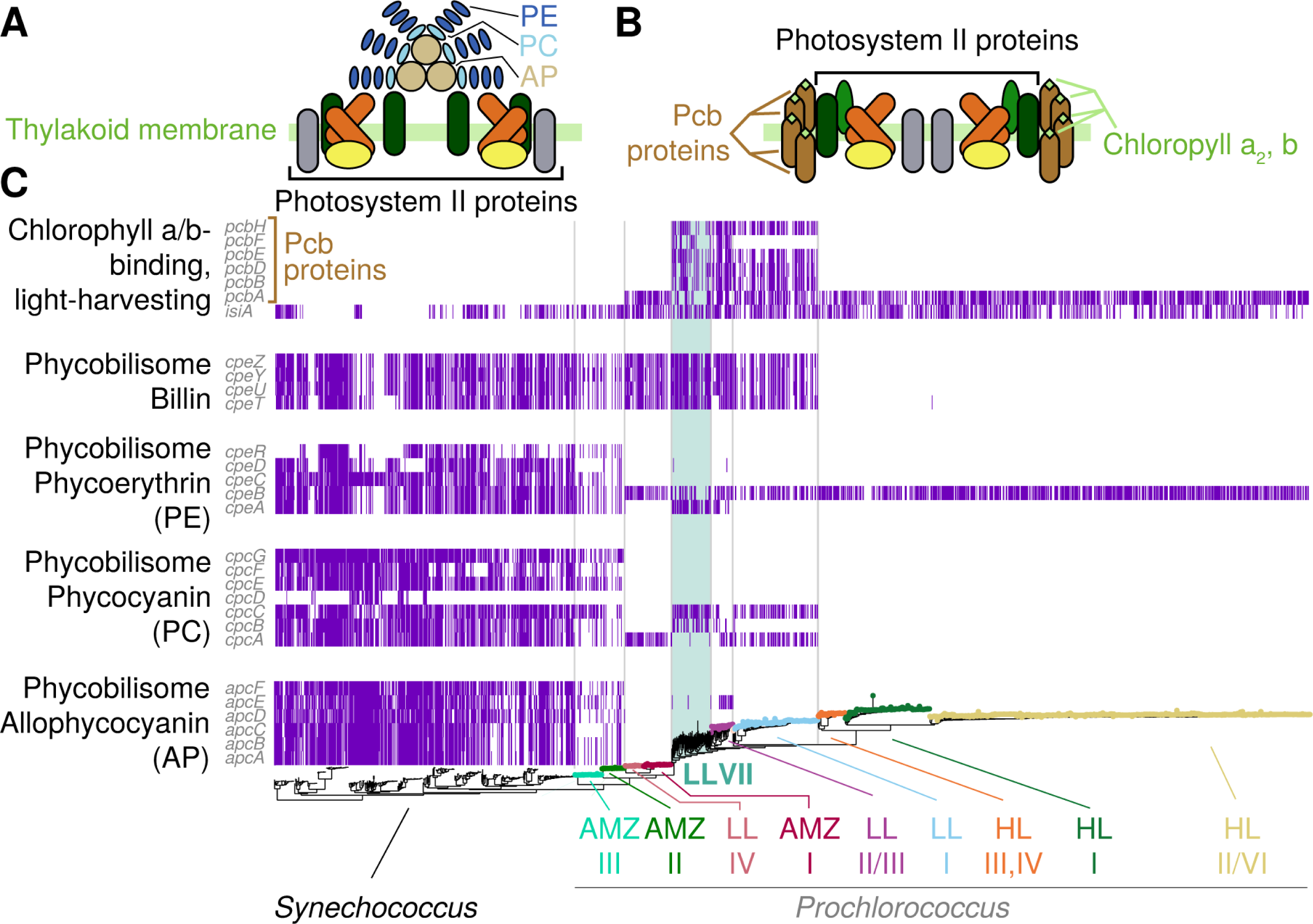
Genomic analysis of photosynthetic pigment biosynthesis and antenna genes across picocyanobacteria. Simplified structure of the phycobillisome (A) and prochlorosome (B) with the major pigments indicated by their abbreviations. (C) Distribution of light-harvesting antennae genes across the phylogeny of marine picocyanobacteria. The tree at the bottom represents the picocyanobacterial phylogeny as in Figure 1, with *Synechococcus* lineages added (see methods). Major lineages are depicted by colored text and corresponding tip point colors, except for *Synechococcus* which has no colored tips, and LLVII which is highlighted with a colored background in the tree and in the matrix. Each row in the matrix represents a single KEGG orthologue (and *isiA*), and colored bars signify the presence of a gene (row) in a genome (column). Vertical lines separate Syn, AMZ-III/II, LLIV/AMZ-I, LLVII, LLI-III, and HL groups. The variation in fine-scale presence/absence patterns observed within clades are most likely due to the inclusion of partial genomes in our database.

Examining the evolution of individual antenna genes reveals a complicated history of divergence and horizontal transfer of multiple genes (Figures S7, S8). In some cases (such as *pcbD*), MIT1223 (LLVIII) shares alleles with LLVII strains, while in others (e.g., *pcbB*), it is closer to LLI strains. Interestingly, MIT1223 never seems to share antenna related alleles with HL strains (Figures S7, S8), suggesting that its “HL-like” characteristics are due to other genetic determinants such as regulation of pigment biosynthesis.

These findings give further support for the important role photophysiology played in the evolution of *Prochlorococcus* (Biller *et al*., 2015), in that major radiations involved significant changes to the configuration of the photosynthetic machinery – more specifically in the light-harvesting apparatus. This began with the complete loss of phycobilisomes in the ancestor of LLIV/AMZI, replaced by the more streamlined *pcb* apparatus (Figure 7, diagram and total number of proteins needed by each antenna type). This transition was capitalized upon by later-diverging lineages, showing remarkable variation in the diversity and copy-number of *pcb* and pigment biosynthesis genes.

### Macroevolutionary implications

Our current understanding of the evolution of *Prochlorococcus* is built upon the notion of an ancestor that once thrived at all depths of the sunlit ocean (Braakman *et al*., 2017). Evolutionary radiations followed, and different lineages emerged with greater relative fitness in shallower waters due to smaller cell size, more streamlined genomes, reconfigured central metabolism, and more optimal elemental compositions. This led to the drawing down of nutrient levels in shallow waters and a displacement of ancestral lineages to deeper waters (Braakman *et al*., 2017). Supporting this view is the distribution of the nitrate reductase gene *narB* in picocyanobacteria, as *narB* is vertically inherited and present in marine *Synechococcus*, LLI, and HL ecotypes, but absent in basal *Prochlorococcus* lineages (Berube *et al*., 2019). This is consistent with the idea that the *Prochlorococcus* ancestor had a broad distribution in the water column and encoded *narB*, which was lost in basal lineages as they were displaced to deeper waters where it was no longer advantageous due to lower light levels and higher levels of other nitrogen sources. Recently, an extension of this model was proposed based on the surprising finding that the basal lineages encode chitinases and that LLIV isolates express functional chitinases and attach to, and use, chitin (Capovilla *et al*., 2023). Under this extension, the *Prochlorococcus* ancestor was particle-associated, and this trait was retained in basal lineages because they still led a facultative particle-associated lifestyle at the base of the euphotic zone, as evident from their enzyme activity and ability to attach to particles and form aggregates. Later radiations, such as the LLVII radiation then fully committed to the planktonic lifestyle and their diversification followed the “top-down” model described above and argued in detail in (Braakman *et al*., 2017).

Our analysis here provides further independent support for the extended model in Capovilla *et al*., 2023. First, it establishes that LLVII isolates are indeed LL-adapted. Second, the massive expansion in *pcb* gene copy number and allelic diversity in the ancestor of the LLVII grade are consistent with the idea that the cells in this radiation switched to a fully planktonic lifestyle. A commitment to the planktonic state, where particle-derived carbon inputs are reduced due to loss of attachment capacities would entail a higher dependence on photosynthetically derived energy and carbon, exerting a strong selective pressure to optimize light-gathering (Capovilla *et al*., 2023). This is supported by the following observations regarding the physiological effect of Pcb proteins in low-light conditions. Different Pcb proteins are known to associate with both photosystems and are the major carriers of chlorophyll in photosystem II, enhancing its light-gathering capacity considerably (Bibby *et al*., 2003). In *Prochlorococcus* strains that carry multiple *pcb* copies, total Pcb levels are higher compared to strains with fewer copies (MacGregor-Chatwin *et al*., 2019), and at least some copies are differentially regulated and are expressed in response to different nutrient limitation conditions. This differential regulation possibly provides specific properties such as variations in trace-element utilization that are beneficial under distinct nutrient limiting conditions (Bibby *et al*., 2003). The distinct physiology and unexpected pigmentation of the LLVIII isolate provide an additional opportunity to investigate the evolutionary journey of *Prochlorococcus* from particle attachment to a free-living lifestyle with greater autotrophic dependence and eventual ability to colonize the upper euphotic zone.

## Conclusions

The four isolates described here help resolve two diverse, largely unexplored regions within the LL-adapted ecotypes of the *Prochlorococcus* collective. The isolates belong to two distinct paraphyletic regions in an area of the *Prochlorococcus* phylogeny with long branch lengths (i.e. the paraphyletic LLVII and LLVIII regions, along with the LLII/III clade). The area is flanked by three monophyletic clades (LLI, AMZI, and LLIV) containing lineages with shorter branch lengths. The inclusion of single cell genomes and metagenomes from field samples provides culture-independent evidence supporting these placements.

A representative from the LLVII grade displayed photophysiology properties and genomic characteristics that were generally intermediate to representatives from the LLIV and HLII clades - two extremes of the *Prochlorococcus* collective life-history spectrum. A representative from the LLVIII grade displayed many photophysiology properties indicative of a LL-adapted strain, however its pigmentation and some genomic characteristics resemble that of a HL-adapted strain, challenging several longstanding paradigms regarding light-adaptation among the *Prochlorococcus* collective. Preliminary work with strain NATL2A (LLI) indicates these anomalies may also extend to members of the LLI clade. Our findings blur the distinction between these historically defined ecotypes and suggest pigmentation and pigment driven photoacclimation are likely unreliable indicators of *Prochlorococcus’* optimal irradiance levels for growth and ecotype designation.

*Prochlorococcus* lineages from regions of the phylogenetic tree with longer branch lengths - and therefore higher genetic divergence from nearby lineages - may reflect divergent evolution among LL-adapted *Prochlorococcus* in which a confluence of environmental variables, including light, has driven substantial diversification from ancestral strains. This is in contrast to the more rapid adaptive radiation events observed within the monophyletic LLI and LLIV clades, which may be a product of finer-scale niche-partitioning among lineages. As the LLVII and LLVIII grades are further examined in the context of evolutionary transitions and the rise of *Prochlorococcus*, the high diversity and long branch lengths contained within these groups may reflect prolonged periods of evolutionary tinkering that occurred during those transitions. Future investigations into the unique photophysiological features we have identified in individual strains, and the genomic and genetic diversity within the grades as a whole, could thus provide important new clues as to how *Prochlorococcus* evolved from its chitin particle-attached ancestors to the planktonic cells that dominate the extant oceans (Braakman *et al*., 2017; Capovilla *et al*., 2023).

## Experimental Procedures

### Isolation and identification of novel *Prochlorococcus* strains

*Prochlorococcus* strains mentioned in this study (Table 1) were isolated from the lower euphotic zone at Station ALOHA in the subtropical North Pacific Ocean (22.75°N 158°W). Strain MIT1223 was isolated from 175 m on September 12, 2012, during the HOE-DYLAN cruise, while all other strains were isolated from 150 m on June 2, 2013, during the HOE-PhoR I cruise. The presence and complexity of *Prochlorococcus* populations were monitored over time by flow cytometry (BD/Cytopeia Influx). The seawater base of all media was sterilized by 0.2 µm filtration followed by autoclaving and prepared in acid-washed autoclaved borosilicate glass or polycarbonate tubes.

MIT1223 was derived from an initial enrichment of unfiltered raw seawater amended with 16 μM NaNO_3_, 1 μM NaH_2_PO_4_.H_2_O, and the trace metal mix used in Pro99 medium (Moore *et al*., 2007) reduced 10-fold. The enrichment was initially maintained for ca. 1 year by serial passage at 22 - 24 °C and ≤ 8 µmol photons m^-2^ s^-1^ on a 14:10 h light:dark cycle, while inorganic nutrient conditions were increased to 80 μM NaNO_3_ and 5 μM NaH_2_PO_4_.H_2_O before acclimating to a modified Pro99 medium (800 μM NH_4_Cl replaced by 800 μM NaNO_3_) and transitioning to higher irradiance - 19 ± 2 µmol photons m^-2^ s^-1^ on a 14:10 h light:dark cycle. Several months later, flow cytometry monitoring revealed the presence of larger picoeukaryote-like cell populations in addition to several *Prochlorococcus* populations. The enrichment was subsequently passed through a 0.8 µm polycarbonate filter (Nucleopore, Whatman/GE) by gentle gravity filtration to remove the larger cells. This enrichment was then maintained by serial passage for another 15 mo before flow cytometry monitoring revealed an apparent unialgal *Prochlorococcus* population, and sequencing revealed the ITS sequence of what we now refer to as strain MIT1223, a member of the paraphyletic LLVIII grade.

Enrichments for *Prochlorococcus* conducted on the HOE-PhoR I cruise in 2013 were prepared as previously described (Cubillos-Ruiz *et al*., 2017; Becker *et al*., 2019). MIT1314 and MIT1327 are derived from an enrichment amended at sea with Pro2 medium nutrients (Moore *et al*., 2007) plus 1 µM thiosulfate, while MIT1300xe, MIT1307xe, and MIT1341xe are derived from an enrichment from the same original 1.0 µm filtered water sample amended with nitrite as the sole exogenous nitrogen source (Table 1). These enrichments and their derivatives from serial passage were maintained at 22 - 25 °C on a 14:10 h light:dark cycle for 3 mo before transitioning to continuous light. Enrichments were maintained at very low irradiances to select for LL-adapted cells, with maximum light exposure of ca. 1 µmol photons m^-2^ s^-1^.

After an initial incubation period of 1 mo, the enrichment that ultimately yielded MIT1314 and MIT1327 was transferred into Pro2 medium plus 1 µM thiosulfate to match the original amendment based on Hawaii surface seawater collected during HOE PhoR I. Two weeks later, a subculture was transitioned to 0.2µm filtered and autoclaved Sargasso seawater amended with Pro99 nutrients (Moore *et al*., 2007). Enrichments were maintained by serial passage in their respective media - MIT1314 derives from the Pro99 line, while MIT1327 derives from the Pro2 line. After 4 mo of serial passage, dilution to extinction in ProMM medium was applied to the Pro2 enrichment, yielding the LLIV clade strain MIT1327 (Cubillos-Ruiz *et al*., 2017). After 5 mo in the laboratory, a subculture of the Pro99 enrichment was moved to higher light (10 ± 2 µmol photons m^-2^ s^-1^) to select for a subset of the *Prochlorococcus* diversity observed by flow cytometry. After six months at this irradiance, ITS screening revealed the HLII clade strain MIT1314.

After an initial incubation period of 7 weeks, the enrichment yielding MIT1300xe, MIT1307xe, and MIT1341xe was maintained by serial passage in Pro99 medium (Moore *et al*., 2007) with a Sargasso seawater base. After 4 mo, replicate subcultures were established and maintained as separate transfer series, with one ultimately resulting in MIT1307xe and the other resulting in MIT1300xe and MIT1341xe. After 1 yr in the laboratory, these enrichments still contained diverse *Prochlorococcus* populations based on flow cytometry and ITS PCR sequencing. Once again, a subculture was transitioned to higher light (10 ± 2 µmol photons m^-2^ s^-1^), and after 1 yr, ITS screening revealed the sequence of what we now refer to as strain MIT1300xe. Flow cytometry monitoring of the two enrichments maintained below 1 µmol photons m^-2^ s^-1^ revealed the presence of LLIV-like *Prochlorococcus* populations and additional *Prochlorococcus* populations with distinct fluorescence signatures. These enrichments were then passed through a 0.8 µm polycarbonate filter (Nucleopore, Whatman/GE) by gentle gravity filtration in an attempt to remove the larger LLIV-like cells. After several months, flow cytometry monitoring revealed an apparent unialgal *Prochlorococcus* population in each, and ITS screening revealed the sequence of what we now refer to as strains MIT1307xe and MIT1341xe.

MIT1314, MIT223, and MIT1327 were purified using a high-throughput dilution to extinction method, and the purity of these isolates was confirmed by flow cytometry and a suite of purity test broths (ProAC, ProMM, and MPTB) as previously described (Berube *et al*., 2014; Cubillos-Ruiz *et al*., 2017; Becker *et al*., 2019). Dilution to extinction purification was repeatedly attempted on the other strains without success.

Initial strain identification was performed by PCR screening of the ITS gene. In brief, 1 ml of culture (ca. 10^6^ to 10^8^ cells) was centrifuged for 15 to 30 min at 16,000 *x g* to form a pellet. The majority of supernatant was removed before centrifugation again at 16,000 *x g* and removal of all residual seawater before resuspending the pellet in 25 to 100 μl Tris-HCl (pH 8.0). Cells were then lysed by boiling at 95 °C for 10 min before centrifugation for 5 min at 16,000 x g at 4 °C to pellet cell debris. The supernatant was removed and used directly in a PCR screen with primers (ITS-F: 5’-CCGAAGTCGTTACTYYAACCC-3’ and ITS-R: 5’-TCATCGCCTCTGTGTGCC-3’), which target the internal transcribed spacer (ITS) region of *Prochlorococcus* as previously described (Rodrigue *et al*., 2009; Berube *et al*., 2015).

### Growth rate experiments

The physiological response of four *Prochlorococcus* isolates (MIT1314, MIT1223, MIT1300xe, and MIT1327) to light availability and temperature was measured in a series of growth experiments. All cultures were maintained in Pro99 medium with a Sargasso surface seawater base in acid-washed autoclaved borosilicate glass tubes.

For light-dependent growth experiments (except NATL2Aax), duplicate tubes of each strain were acclimated to target light levels under continuous light and monitored weekly using a light meter (LI-COR LI-250A) connected to a spherical micro quantum sensor (Walz US-SQS/L). 95% of irradiance measurements were within 10% of the target light level for strains MIT1223, MIT1300xe, and MIT1327 and within 16% of the target light level for MIT1314 (Table S1;Figure S9). Temperature was monitored daily and held stable at 24 ± 1 °C. Balanced growth was confirmed by examining growth rates over time at each condition. Attempts were made to acclimate each strain to increasingly higher and lower light intensities until reproducible growth could not be achieved. Growth was monitored daily by flow cytometry (see below) and bulk chlorophyll fluorescence (10AU model, Turner Designs) for 4 - 6 days. Bulk chlorophyll fluorescence readings were blank subtracted using the mean of triplicate medium blanks, and any readings corresponding to lag phase or early onset of stationary phase were identified by eye and removed prior to calculating mean growth rates (Table 2A). For light-dependent growth experiments with NATL2Aax, triplicate tubes were acclimated to 4 target light levels (10, 20, 45, and 96 ± 0.5 µmol photons m^-2^ s^-1^) on a 14:10 h light:dark cycle. Growth was monitored daily by flow cytometry (see below) and bulk chlorophyll fluorescence (10AU model, Turner Designs) for 8 - 16 days, and mean growth rates were calculated from blank subtracted fluorescence readings taken in exponential phase.

For temperature-dependent growth experiments, duplicate tubes of each strain were acclimated to specific temperatures (± 0.5°C) and grown at (76 ± 1 µmol photons m^-2^ s^-1^ for MIT1314; 20 ± 1 µmol photons m^-2^ s^-1^ for MIT1223, MIT1300xe, and MIT1327) on a 14:10 h light:dark cycle. Acclimation was defined as described above, and attempts were made to acclimate each strain to increasingly higher and lower temperatures until reproducible growth could not be achieved. Growth was monitored daily, and mean growth rates were calculated as described above (Table 2B).

### Flow cytometry and fast repetition rate fluorometry

Cell concentrations were determined using a Guava Technologies easyCyte 12HT flow cytometer (EMD Millipore). Samples were diluted in sterilized Sargasso seawater to ensure <500 cells µl^-1^ to avoid coincidence counting and were run with only the blue (488 nm) excitation laser enabled for maximum power. Technical duplicate *Prochlorococcus* populations were resolved for each biological sample based on their red (695/50 nm) emission parameters for enumeration. A fluorescent bead reagent (easyCheck Beads, EMD Millipore) was run daily as a reference, and to verify instrument performance. Median forward angle scatter, chlorophyll (695/50 nm), and yellow (583/26 nm) fluorescence per cell were determined for each population.

The maximum quantum efficiency of photochemistry in PSII (Fv/Fm) and the functional absorption cross-section of PSII (σPSII) were measured using fast repetition rate fluorometry (FRRF) on a FIRe fluorometer instrument (Satlantic) as previously described (Biller, Coe, *et al*., 2018). Samples were stored in the dark for 15 min prior to loading into the instrument and Pro99 medium was used for blank measurements.

### Absorption spectra and pigment measurements

*In-vivo* absorption spectra were obtained using a Beckman DU 800 spectrophotometer (Beckman Coulter Inc.) in absorbance scan mode from 400 to 700 nm at 1.0 nm intervals and a scan speed of 1200 nm/min and Pro99 medium was used for blank measurements.

For pigment sampling, duplicate *Prochlorococcus* cultures, as well as technical duplicates from each biological replicate, were filtered using vacuum filtration (ca. -200 mm Hg) onto 47 mm diameter 0.2 µm hydrophilic Durapore filters (Millipore). Samples were immediately flash-frozen and stored in liquid nitrogen (-196 °C) until processing. Pigments were extracted using a modified Bligh and Dyer protocol (Popendorf *et al*., 2013) with DNP-PE-C_16:0_/C_16:0_-DAG (2,4-dinitrophenyl phosphatidylethanolamine diacylglycerol; Avanti Polar Lipids, Inc., Alabaster, AL) used as an internal standard. Filter blanks and *Prochlorococcus* growth media blanks were extracted and analyzed alongside samples. The total lipid extract was analyzed by reverse-phase high-performance liquid chromatography (HPLC) mass spectrometry (MS) on an Agilent 1200 HPLC coupled to a Thermo Fisher Exactive Plus Orbitrap high-resolution mass spectrometer (ThermoFisher Scientific, Waltham, MA, USA). HPLC and MS conditions were as previously described (Hummel *et al*., 2011; Collins *et al*., 2016). In brief: 20 µl were injected onto a C8 Xbridge HPLC column (particle size 5 µm, length 150 mm, width 2.1 mm; Waters Corp., Milford, MA, USA). Lipids were eluted at a flow rate of 0.4 ml min^-1^ using the following gradient with eluent A (water with 1% 1M ammonium acetate and 0.1% acetic acid) and eluent B (70% acetonitrile, 30% isopropanol with 1% 1M ammonium acetate and 0.1% acetic acid): 45% A was held for 1 min, from 45% A to 35% A in 4 min, from 25% A to 11% A in 8 min, from 11% A to 1% A in 3 min with an isocratic hold until 30 min. Finally, the column was equilibrated with 45% A for 10 min. ESI source settings were: Spray voltage, 4.5 kV (+), 3.0 kV (-); capillary temperature, 150 °C; sheath gas and auxiliary gas, both 21 (arbitrary units); heated ESI probe temperature, 350 °C. Mass data were collected in full scan while alternating between positive and negative ion modes. Pigments were identified using retention time, accurate molecular mass, and isotope pattern matching of proposed sum formulas in full-scan mode and tandem MS (MS^2^) fragment spectra of representative compounds. For each MS full scan, up to three MS^2^ experiments targeted the most abundant ions with N2 as collision gas. The scan range for all modes was 100-1500 *m/z*. The mass spectrometer was set to a resolving power of 140,000 (FWHM at *m/z* 200), leading to an observed resolution of 75,100 at *m/z* 875.5505 of our internal standard, DNP-PE. Exact mass calibration was performed by weekly infusing a tune mixture. Additionally, every spectrum was corrected using a lock mass, providing real-time calibrations. To validate the accuracy and reliability quantification, quality control samples of known composition spiked with lipid standards were interspersed with the samples as described previously (Collins *et al*., 2016). Pigment abundances were corrected for the relative response of commercially available standards. The abundances of chlorophylls and their associated compounds were corrected for the response of a chlorophyll *a* standard and carotenoid pigments using a *β*-carotene standard. All standards were purchased from Sigma Aldrich (St. Louis, MO, USA). Individual response factors were obtained from external standard curves by triplicate injection of a series of standard mixtures ranging from 0.15 to 40 pmol on column per standard. Our method’s use of external standards was validated in a study that compared lipid quantitation against internal, isotope-labeled standards (Becker *et al*., 2018). Data were corrected for differences in extraction efficiency using the recovery of the DNP-PE internal standard.

### Phylogenetic analysis

Maximum likelihood trees were reconstructed using FastTree (Price *et al*., 2010) from 44 concatenated single-copy core proteins that have conserved local synteny (see below) in 1691 non-redundant Picocyanobacterial genomes (1201 *Prochlorococcus* and 490 *Synechococcus*). 4512 genomes matching either *Prochlorococcus* or *Synechococcus* in their description were downloaded from NCBI and IMG on December 7 2022. To filter out very closely related sequences, we used FastANI (Jain *et al*., 2018) to calculate all pairwise whole-genome average nucleotide identity (ANI) values. From these results a network was created where nodes represent genomes and links represent >99.99% identity over 95% of the longer genome in the pair. This network had 1961 connected components, and the longest genome from each component was picked as the representative, resulting in the final genome set used in downstream analysis. Gene calling was performed using Pyrodigal v2.0.2 (Larralde, 2022) and protein sequences were clustered using MMseqs2 v14-7e284 (Steinegger and Söding, 2017) based on 50% identity and 80% coverage. Initial protein homology clusters were then split based on their genomic surroundings into groups that share at least 3 of the 10 protein families in their immediate genomic vicinity. The final protein clusters therefore share both homology and local synteny. We found 44 final clusters that are present in >50% of analyzed genomes, and calculated the protein alignment of each protein cluster using mafft v7.310 (Katoh and Standley, 2013). We then transformed the protein alignments to nucleotide alignments and concatenated all individual alignments to one long alignment that was used for phylogenetic inference with FastTree v2.1.10 (Price *et al*., 2010).

### Genome sequencing and analysis

Closed genome sequences of MIT1223, MIT1300xe, MIT1307xe, and MIT1341xe were obtained as previously described for MIT1314 (Becker *et al*., 2019). In brief, genomic DNA was extracted from concentrated cells using the MasterPure complete DNA and RNA purification kit (Epicentre) and sequenced using P6 chemistry on a RS II instrument (Pacific Biosystems). Library construction and sequencing were performed at the University of Massachusetts Medical School Deep Sequencing and Molecular Biology Core Laboratory. The average insert range for libraries constructed using a standard SMRTbell Template Prep Kit 1.0 and Sequencing Primer v3 (Pacific Biosciences) was 21 – 28 kb. Reads were assembled *de novo* into a single contig for each strain using the RS_HGAP_Assembly.2 protocol within the SMRT Analysis software (v2.0; Pacific Biosciences) before circularization using the Geneious sequence analysis package (V7.1, Biomatters) and polishing using Quiver and the RS_Resequencing.1 protocol within the SMRT Portal software. The average coverage for each assembly ranged from 73X to 335X (Table 3). The draft genome of MIT1327 was obtained as previously described (Cubillos-Ruiz *et al*., 2017) and consists of 29 scaffolds with a N50 of 328.4 kbp and an average coverage of 204X (Table 3). All genomes have been deposited in GenBank at the National Center for Biotechnology Information (NCBI) and the Joint Genome Institute’s Integrated Microbial Genomes (IMG) system (see Table 3 for IDs) and annotated using the IMG Annotation Pipeline version 4 (Markowitz *et al*., 2013; Huntemann *et al*., 2016). Sequenced isolates were included in the genome collection described above.

The proteomes of all genomes were annotated using the eggnog v6 database (Hernández-Plaza *et al*., 2022, 6). Input proteins were competitively aligned to the database using MMseqs2 with a minimum identity of 30% and 50% coverage. The annotations of the best annotated hit of each input protein was then transferred to the input protein. For the analysis in figure 7, antenna proteins in the “Photosynthesis Proteins” kegg pathway (ko00194) were extracted based on their eggnog-derived KO numbers, and their presence/absence in genomes was tabulated and plotted on the phylogenetic tree. For each KO the phylogenetic tree was built by aligning KO members using mafft and building a tree using FastTree as above. Gene trees were rooted using the minimal ancestor deviation criterion (Tria *et al*., 2017).

## Acknowledgements

We thank the captains, crews, and research teams of the HOE-DYLAN and HOE-PhoR I research cruises, particularly chief scientists Sam Wilson and Karin Björkman, for facilitating *Prochlorococcus* enrichments. We also thank Kelsey Perry for assistance with pigment extractions. This work was supported in part by grants from the National Science Foundation (OCE-1153588 and DBI-0424599 to S.W.C.), the Simons Foundation (Life Sciences Project Award ID 337262, S.W.C; SCOPE Award ID 329108, S.W.C), and the Gordon and Betty Moore Foundation (Grant IDs GBMF495 and GBMF4511 to S.W.C.). This paper is a contribution from the Simons Collaboration on Ocean Processes and Ecology (SCOPE) and the NSF Center for Microbial Oceanography: Research and Education (C-MORE).

---

**Figure S1.**
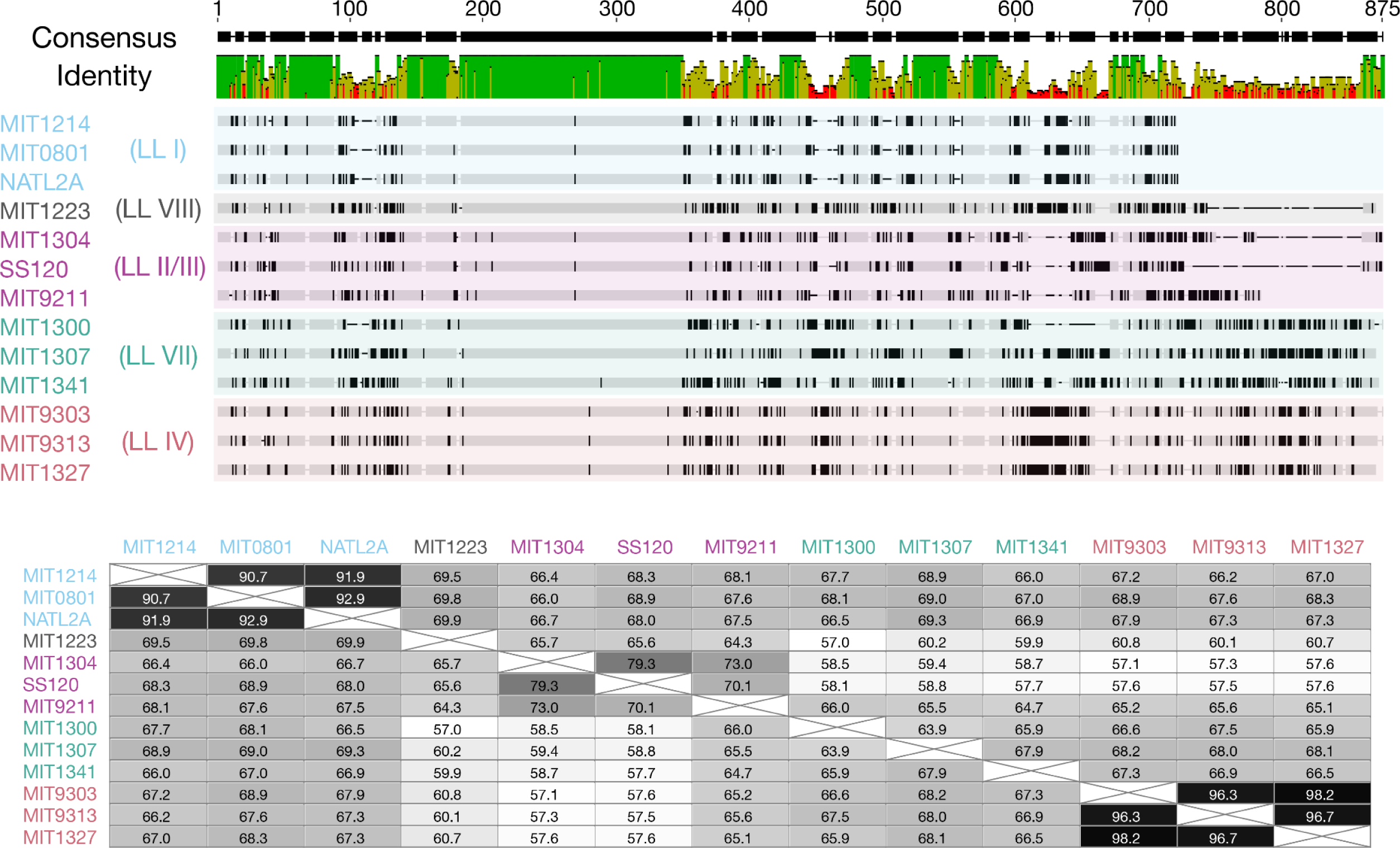
MAFFT (V7.54; (Katoh and Standley, 2013) alignment (top) of internal transcribed spacer (ITS) regions derived from *Prochlorococcus* isolates representing established clades (LLI, LLII/III, LLIV) and novel grades (LLVII, LLVIII) created using default settings in the Geneious software package (V10.2.6; (Kearse *et al*., 2012). A percent identity matrix for the alignment is also included (bottom).

**Figure S2.**
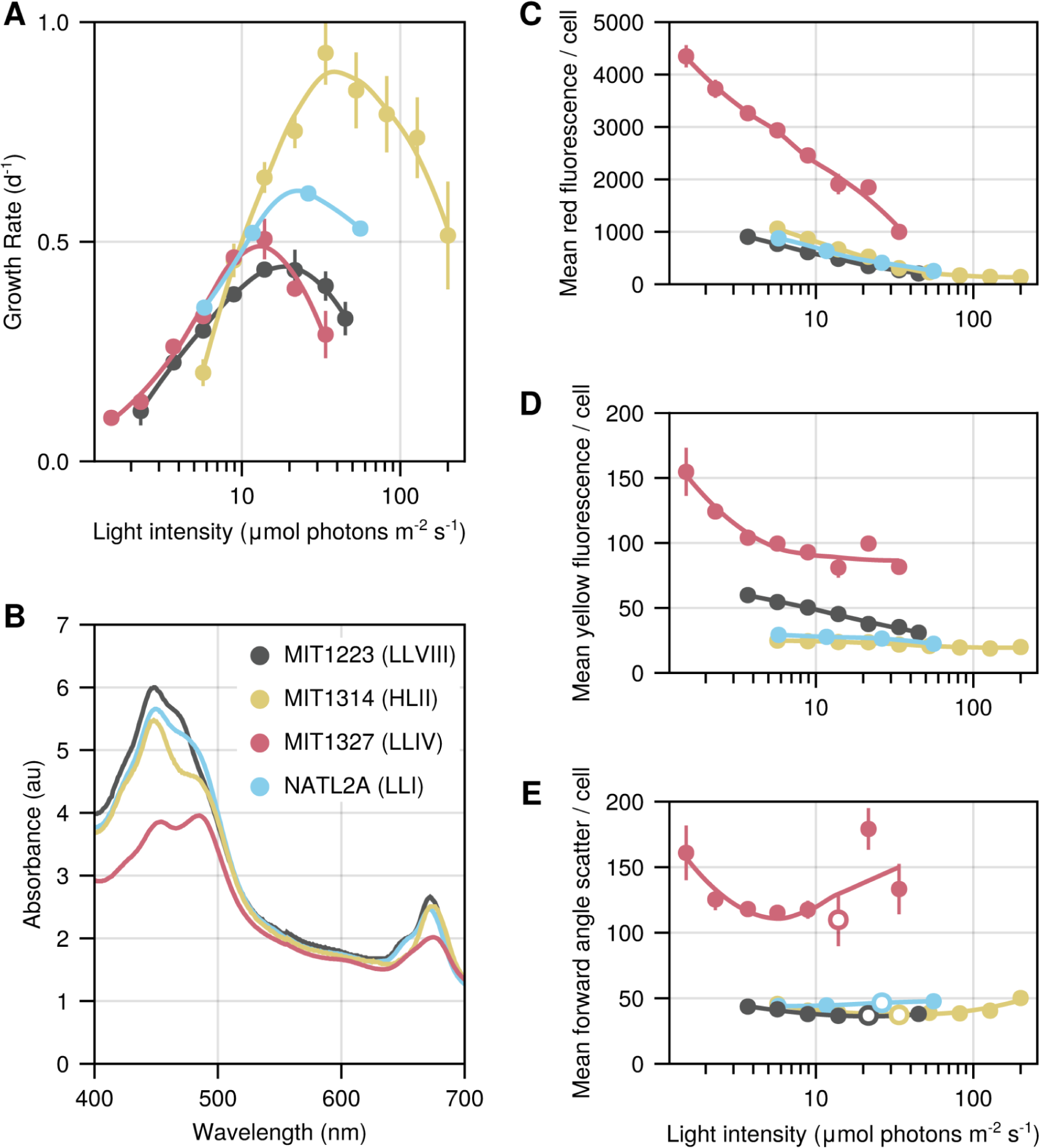
Photophysiology comparison with NATL2A (LLI clade). Light-dependent growth rates (A), *in vivo* absorption spectra for log phase cultures of different strains growing at 5.7 - 5.8 μmol photons m^-2^ s^-1^ (B), and flow cytometry parameters red fluorescence (C), yellow fluorescence (D), forward scatter (E) for NATL2A - a well studied LL-adapted strain of *Prochlorococcus* (Scanlan *et al*., 1996; Biller *et al*., 2015; Berube *et al*., 2019) - for comparison with the novel strains reported in this study. Open circles denote the light level at which maximum growth rate was achieved for each strain (E) Data for MIT1314, MIT1223, and MIT1327 are reproduced from figures 2, 4, and 5.

**Figure S3.**
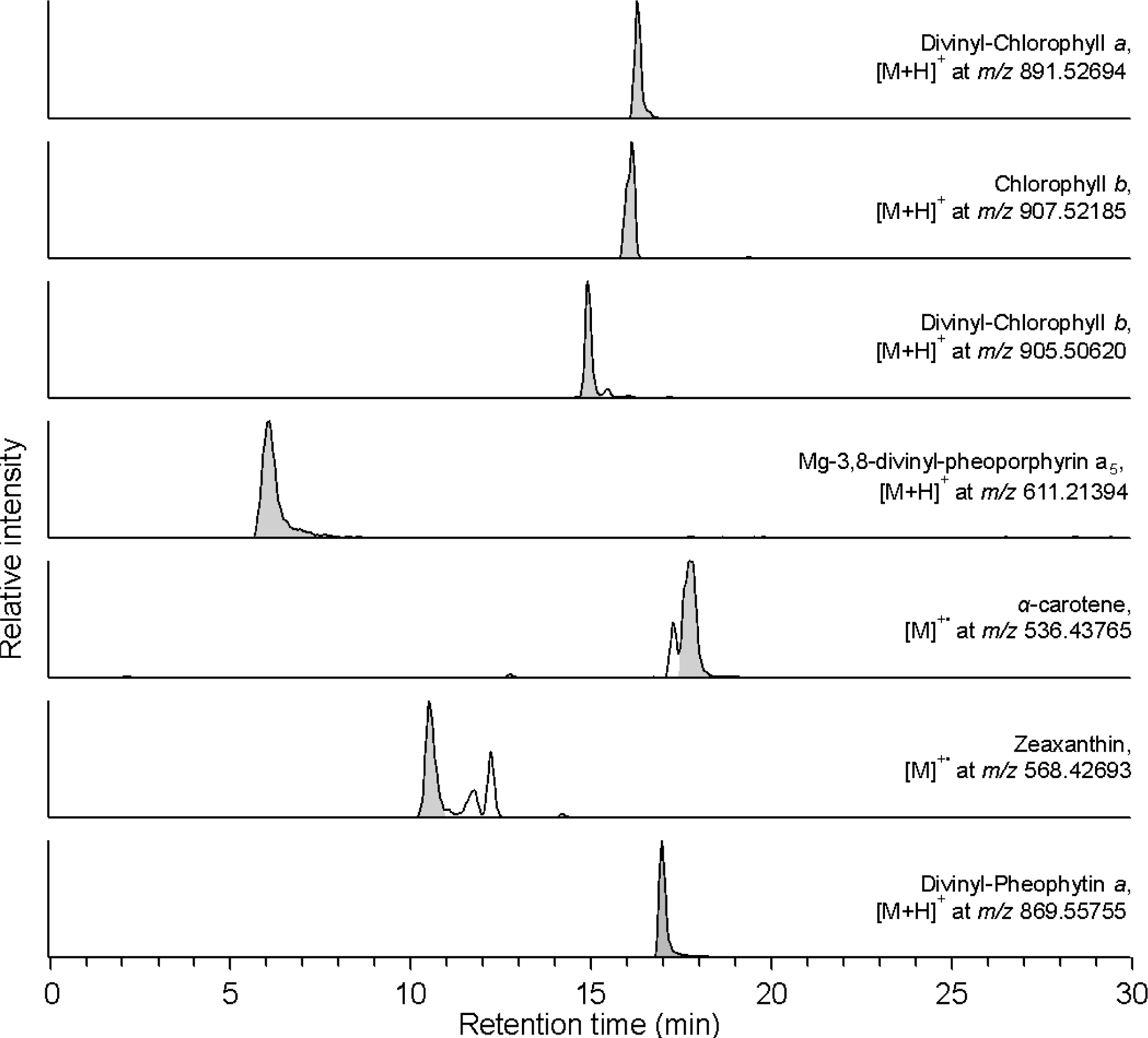
HPLC-Orbitrap-MS chromatograms shown as extracted ion chromatograms (EICs) of identified pigments in *Prochlorococcus* MIT1314 (HLII clade) acclimated to an irradiance of 21.6 µmol photons m^-2^ s^-1^.

**Figure S4.**
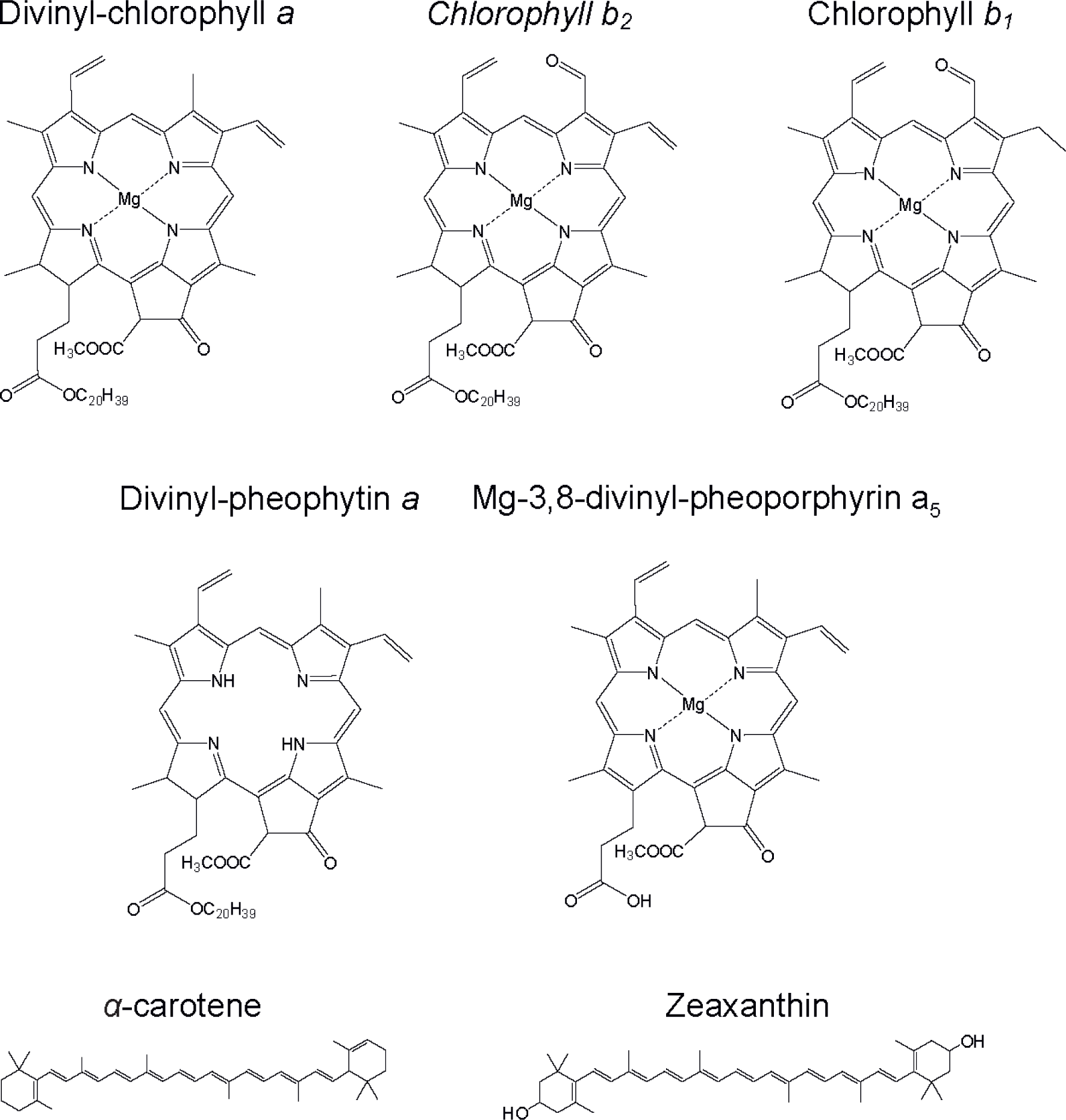
Molecular structures of pigments identified in *Prochlorococcus* strains MIT1327, MIT1300xe, MIT1223, and MIT1314.

**Figure S5.**
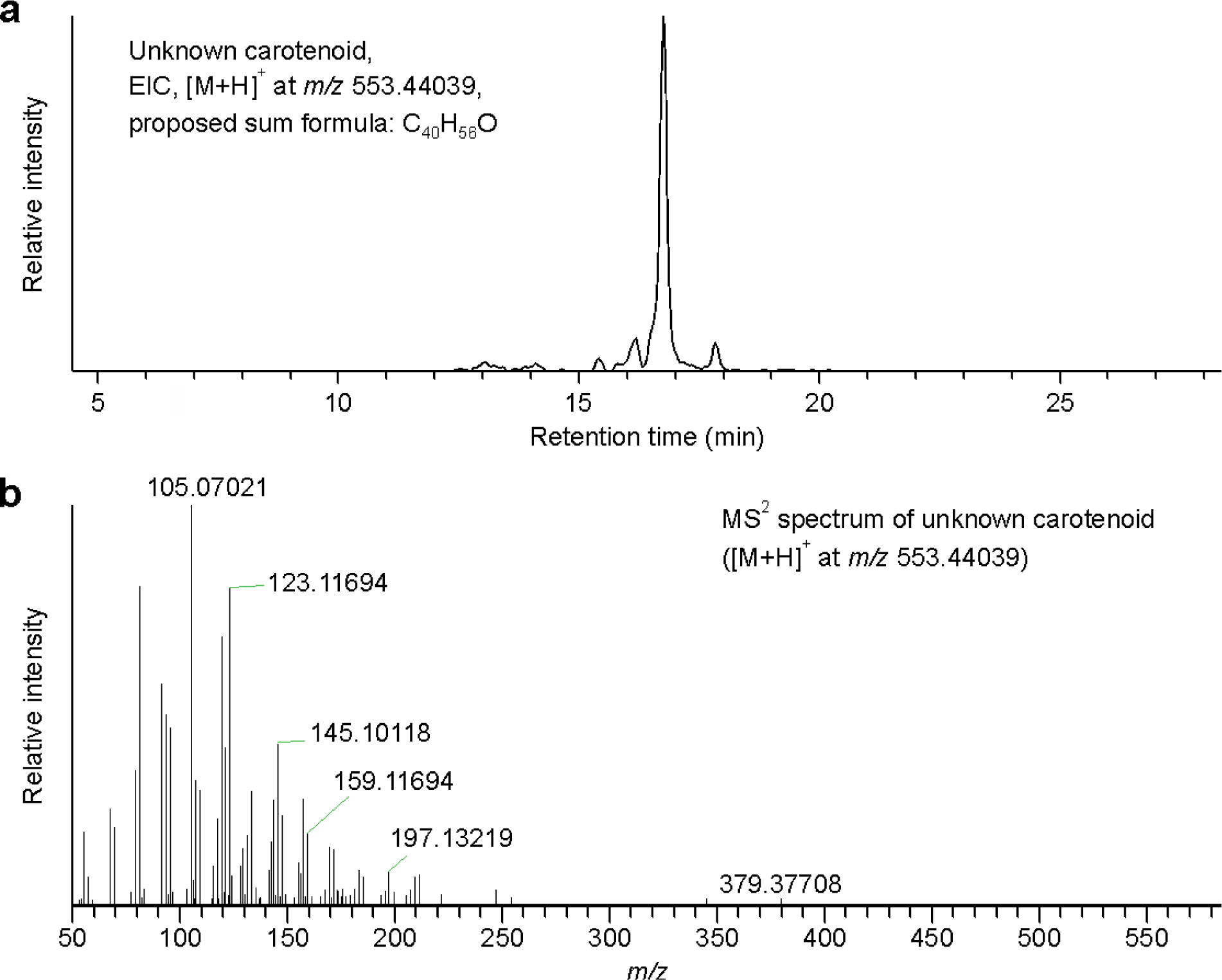
HPLC-Orbitrap-MS chromatogram shown as extracted ion chromatogram (EIC) of an unknown pigment ([M+H]^+^ at *m/z* 553.44039) in *Prochlorococcus* MIT1314 acclimated to an irradiance of 21.6 µmol photons m^-2^ s^-1^ (A). The proposed sum formula (C_40_H_56_O) based on the accurate mass in full scan is also shown. MS^2^ spectrum of the unknown carotenoid ([M+H]^+^ at *m/z* 553.44039) (B). We observed a dominant fragment ion at *m/z* 123.117 in the MS^2^ spectrum, which is typically associated with *α*-carotene (Rivera *et al*., 2014) as the fragment represents the ε ring. The second dominant fragment at *m/z* 105 is usually associated with epoxy xanthophylls (Rivera *et al*., 2014).

**Figure S6.**
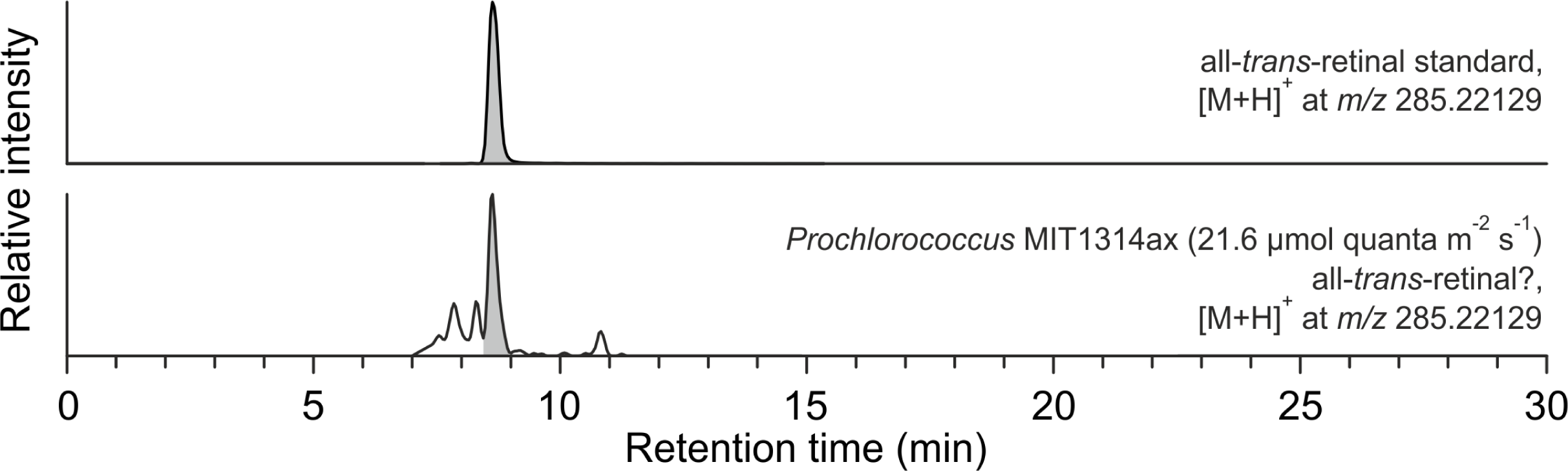
HPLC-Orbitrap-MS chromatogram shown as extracted ion chromatogram (EIC) of an all-*trans*-retinal standard ([M+H]^+^ at *m/z* 285.22129) and an EIC of the same mass in *Prochlorococcus* MIT1314 acclimated to an irradiance of 21.6 µmol photons m^-2^ s^-1^. The match of the accurate mass together with the retention time indicates the presence of all-*trans*-retinal in *Prochlorococcus*, but further identification is needed to confirm this identification. Due to its low abundance, we were not able to obtain MS^2^ spectra of the putative retinal

**Supplementary Figure 7:**
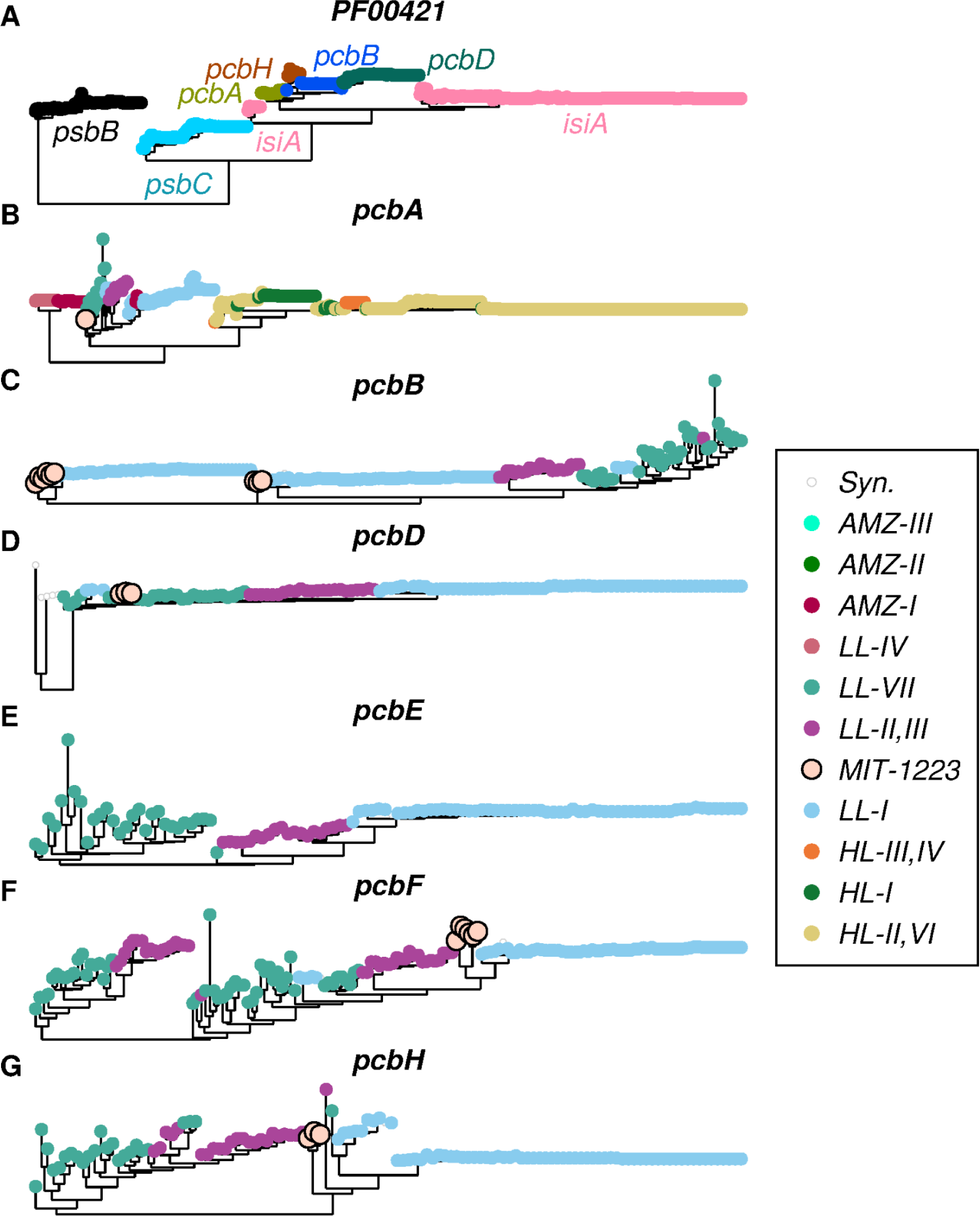
Trees of picocyanobacterial genes that contain the pfam PF00421 (Photosystem II protein) domain. (A) Sequences were extracted based on the presence of the PF00421 domain. Colors represent assigned annotation, as used in Figure 8. pcbE/F proteins only contain weak hits to PF00421, and were excluded from the tree. For psbB/C, 100 sequences were chosen at random, as they are all monophyletic. (B-G) trees of individual genes with tip colors depicting ecotype as shown in the legend. (B) pcbA mostly fits the species phylogeny and shows very different alleles for Basal (LL-IV, and AMZ-I), LL, and HL clades. (C) pcbB is found in LL only. MIT1223 together with LL-I. LL-I also had a duplication, which MIT1223 lacks. (D) MIT1223 embedded in LL-VII. (E) agrees with the species tree. (F) MIT1223 closest to LL-I. (G) MIT1223 has a divergent allele, dissimilar to both LL-VII and LL-I.

**Supplementary Figure 8:**
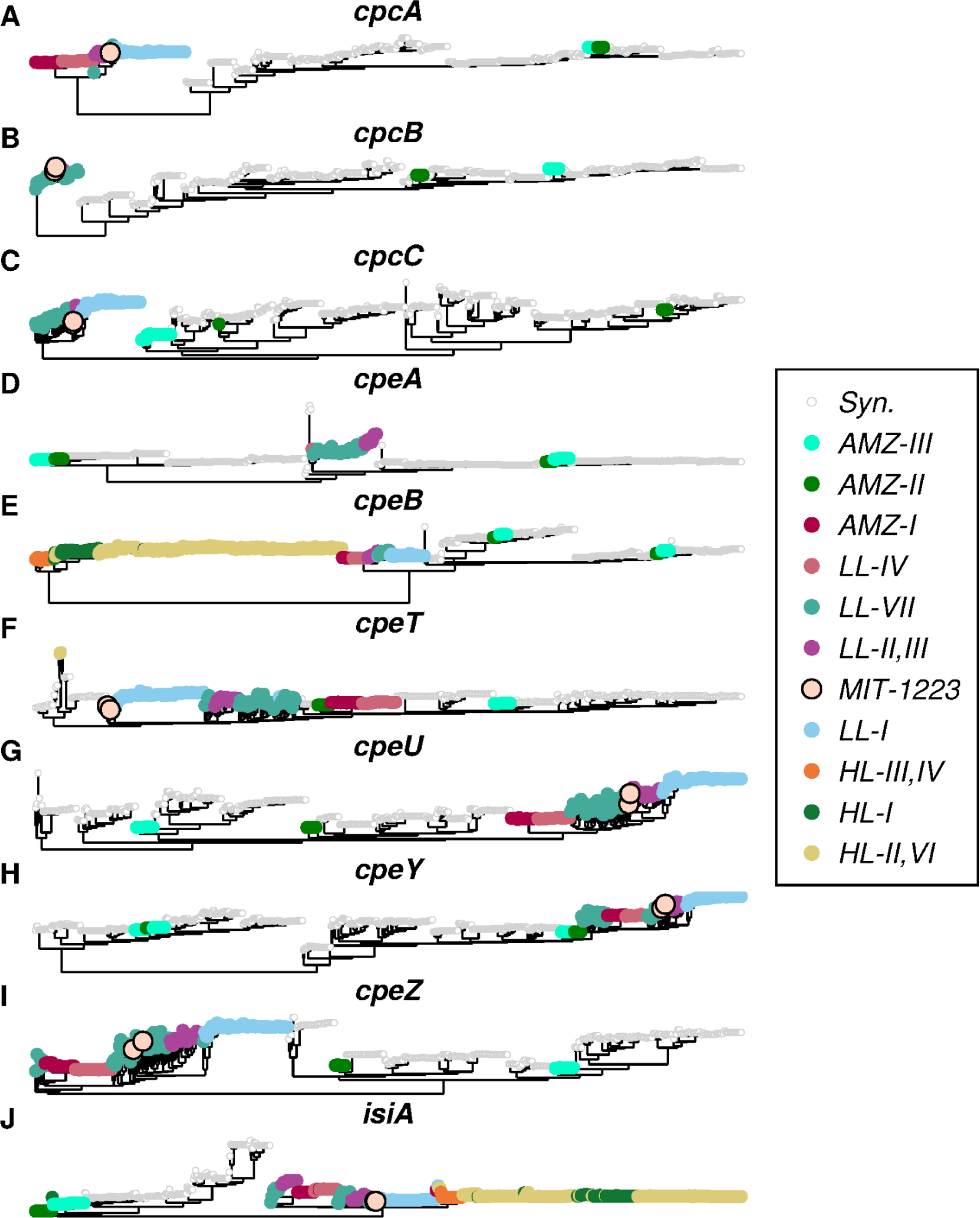
Trees of phycobilisome associated genes. Tip colors represent ecotype as denoted in the legend. (A) *Prochlorococcus* allele is completely different from *Synechococcus*, except for AMZ-III, and II, which share the *Synechococcus* allele. (B) MIT1223 embedded in LL-VII. *Prochlorococcus* allele is completely different from *Synechococcus*, except for AMZ-III, and II, which share the *Synechococcus* allele. (C) MIT-1223 has a divergent allele, dissimilar to both LL-VII and LL-I. (D) Duplication in *Synechococcus*. AMZ-II and III acquired both copies. LL-VII and LL-II,III have a different allele that diverged from the ancestor of *Synechococcus* genes before their duplication. (E) HL allele is completely different from rest. Basal *Prochlorococcus* (AMZ-I, LL-IV, LL-II.III, LL-I, and LL-VII) have a different allele than *Synechococcus*, which has a duplication that is shared with AMZ-II and III. (F) MIT1223 has a divergent allele, dissimilar to both LL-VII and LL-I. Basal *Prochlorococcus* (AMZ-I to AMZ-III, and LL-IV) are embedded in *Synechococcus* diversity. (G) *Prochlorococcus* portion agrees with the species tree. AMZ-II and AMZ-III each have a different *Synechococcus* allele. (H) Duplication in *Synechococcus*. AMZ-II, and II have both *Synechococcus* alleles. *Prochlorococcus* diversity agrees with the species tree, and branches off from one of the *Synechococcus* alleles. (I) Three clades in *Synechococcus*. AMZ-II, and AMZ-III each have a different *Synechococcus* allele. MIT1223 embedded in LL-VII diversity. (J) *Prochlorococcus* completely separated from *Synechococcus*. Complex history in *Pro*. MIT1223 and some LL-VII share the LL-I allele.

**Figure S9.**
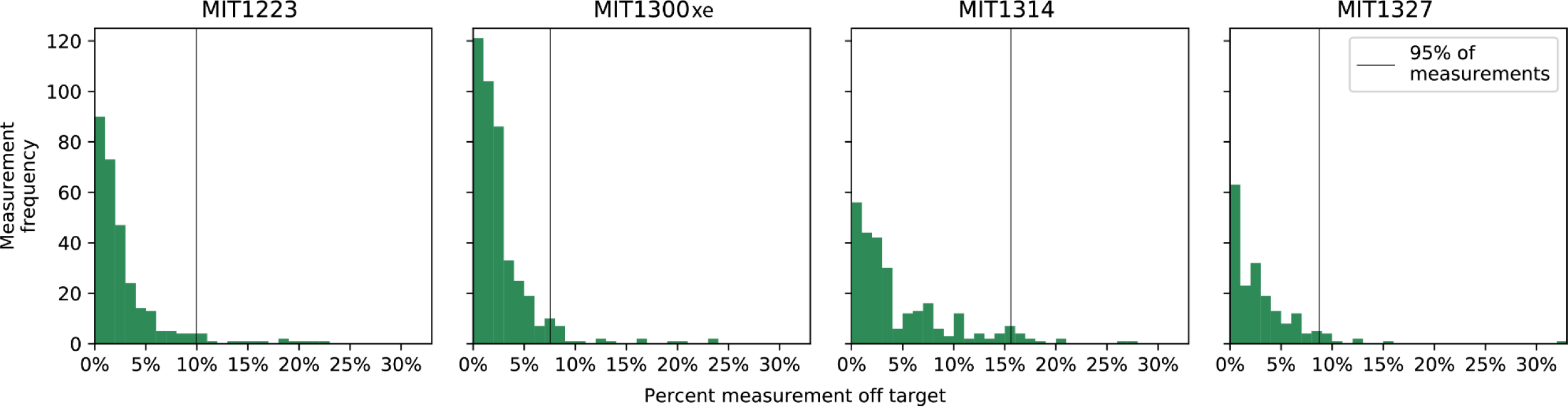
Histograms showing percent deviation from target light level for irradiance measurements taken over the course of light-dependent growth experiments.

**Table S1.** Summary statistics for light measurements (µmol photons m^-2^ s^-1^) taken over the course of the light-dependent growth rate experiments. Measurements were taken weekly and minor adjustments were made to maintain target levels. Two of the target light levels listed above (40.0 and 73.0 µmol photons m^-2^ s^-1^) were used for acclimation purposes – no samples were taken at these levels.

| Target light level | Median light level | Mean light level | Standard deviation | Minimum value | Maximum value | CV | Mean % of target | 2.5th percentile | 97.5th percentile |
| --- | --- | --- | --- | --- | --- | --- | --- | --- | --- |
| 1.5 | 1.5 | 1.5 | 0.1 | 1.0 | 1.6 | 5.5 | 98.0 | 1.3 | 1.6 |
| 2.3 | 2.3 | 2.3 | 0.1 | 1.8 | 2.8 | 5.3 | 98.8 | 1.9 | 2.5 |
| 3.7 | 3.7 | 3.7 | 0.2 | 2.9 | 4.5 | 5.7 | 99.6 | 3.2 | 4.0 |
| 5.7 | 5.7 | 5.7 | 0.3 | 4.8 | 6.5 | 4.7 | 100.0 | 5.2 | 6.3 |
| 8.9 | 8.8 | 8.7 | 0.4 | 7.3 | 9.8 | 4.2 | 98.2 | 7.7 | 9.3 |
| 13.9 | 13.8 | 13.7 | 0.6 | 11.6 | 16.2 | 4.3 | 98.8 | 12.3 | 14.7 |
| 21.6 | 21.4 | 21.4 | 1.1 | 16.6 | 26.0 | 5.3 | 99.0 | 18.5 | 23.2 |
| 33.8 | 33.7 | 33.5 | 1.6 | 27.0 | 39.1 | 4.7 | 99.2 | 29.4 | 36.4 |
| 40.0 | 39.6 | 39.2 | 0.9 | 38.0 | 40.1 | 2.2 | 97.9 | 38.0 | 40.1 |
| 45.0 | 44.5 | 44.2 | 2.0 | 38.6 | 48.0 | 4.5 | 98.3 | 39.2 | 47.1 |
| 52.7 | 52.0 | 51.9 | 2.6 | 44.0 | 66.5 | 4.9 | 98.4 | 46.9 | 55.7 |
| 65.0 | 63.2 | 63.6 | 3.0 | 57.0 | 76.0 | 4.8 | 97.8 | 57.6 | 69.8 |
| 73.0 | 75.3 | 75.1 | 3.3 | 70.0 | 81.0 | 4.4 | 102.8 | 70.2 | 80.8 |
| 82.2 | 82.0 | 81.5 | 5.7 | 65.7 | 104.4 | 7.0 | 99.2 | 69.2 | 90.3 |
| 128.0 | 129.5 | 128.0 | 8.7 | 108.0 | 151.2 | 6.8 | 100.0 | 109.5 | 149.2 |
| 200.0 | 192.0 | 191.5 | 10.2 | 166.0 | 212.0 | 5.3 | 95.7 | 170.2 | 210.9 |

